# Genomic properties representing plant sex chromosome evolution interpreted with genome language models

**DOI:** 10.64898/2026.05.25.727704

**Authors:** Takashi Akagi, Hikaru Matsuoka, Jun Takayama, Gen Tamiya

## Abstract

Plants have repeatedly evolved chromosomal sex-determining systems from hermaphroditic ancestors, providing a powerful natural framework for studying convergent evolution. However, sex chromosomes undergo extensive structural divergence, degeneration, and repeat accumulation, making direct comparisons across distant lineages difficult. Here, we apply a genome language model (gLM), which encodes genomic sequences into high-dimensional representations of their contextual properties, to independently evolved sex chromosomes from the distantly related genera *Silene* and *Humulus*. Without relying on sequence alignment or gene orthology, we identify convergent genomic signatures shared among plant Y chromosomes, including elevated GC content and depletion of specific trinucleotide motifs. Directionality analyses of latent genomic vectors further reveal common evolutionary trajectories associated with recombination suppression and Y chromosome differentiation. These properties differ from those observed in animal sex chromosomes, suggesting lineage-specific modes of sex chromosome evolution in plants. Our results demonstrate that genome language models can transform structurally incomparable chromosomes into quantitatively comparable evolutionary entities, allowing the interpretation of common genomic principles underlying convergent evolution across deeply diverged plant lineages.

## Introduction

Plants and animals differ in their sexual systems. In animals separate sexes are common, and chromosomal sex-determining systems are widespread. In mammals, for example, individuals’ sexes are almost always controlled by nearly identical sex-determining systems involving the actively male-determining *Sry* gene^1–2^. In contrast, hermaphrodite or monoecious species predominate in flowering plants, and are ancestral to many different lineages that have independently evolved chromosomal sex-determining systems that control development of separate male and female individuals, which is termed dioecy^3–4^. Insights into plant sex-determining and sex chromosome evolution have benefitted from empirical and theoretical studies^5–11^, the former especially through cytogenetic and genetic studies, many using *Silene latifolia* and its close relatives in the family Caryophyllaceae^12–18^. Recently, advances in genome sequencing have yielded complete assemblies of the sex chromosome of several dioecious Angiosperm species, such as in persimmon^19–21^, garden asparagus^22^; kiwifruit^23–24^, *Salix* specis^25–27^; spinach^28–29^, *Rumex hastatulus*^30^, *Silene* species^31–33^, hops^34^, and their likely outgroup *Amborella*^35^. Analyses of these genomes have confirmed the conclusion from cytogenetic and early genetic studies that the extent of Y chromosomes differentiation (including heterochromatinization) compared with their ancestral states, represented by the X chromosomes of the same species, differs greatly among species^5^. The sex chromosome pairs of some species include large non-recombining regions. In species with male heterogamety, these “male-specific regions of the Y” (MSY) can differ greatly in size and degree of heterochromatinization from their X counterpart (X-Y heteromorphism) and genetic degeneration, with loss of genes present on the X. In other species, most of the sex chromosome pair continues to recombine with the X, and remains pseudoautosomal (PAR), with no extensive MSY, largely similar X and Y gene contents and orders (synteny). Although heteromorphism can arise in other ways, such as fusions between a sex chromosome and an autosome, and this can explain the lack of a clear relationship with the evolutionary ages of plant sex chromosome systems^36^, cessation of recombination is of great importance, as it initiates several of the processes that can explain their consistently observed size changes (a major category of heteromorphism), losses of genes (sometime profound losses) and heterochromatinization, including repetitive sequence accumulation. Certain rearrangements can also create differences in morphology, including pericentric inversions, but these too are unlikely genomic events except in regions that recombine rarely or not at all.

To date, fully resolved sequences of heteromorphic Y chromosomes have been reported only for the genera *Silene* (*S. latifolia* and *S. dioica*)^32^ and *Humulus* (*H. lupulus* and *H. japonicus*)^34^. Although these genera diverged more than 100 million years ago^37–39^, their Y chromosomes evolved over short evolutionary periods (*dS* < 0.25). Therefore, a comparative analysis of these independently evolved Y chromosomes may reveal features common to evolution of such sex chromosomes in plants.

The evolution of heteromorphic sex chromosomes in plants or animals involves the establishment of large non-recombining regions. Under male heterogamety (which we consider from now onwards, as most concepts apply to female heterogamety with the sexes appropriately changes), this leads to a low effective population size (*N*_e_) compared with that of the X chromosome (because, with a 1:1 sex ratio, the population includes three times as many of the haplotype for the X-linked region, compared with the Y-linked region). Several effects collectively further reduce the Y effective population size, including Hill-Robertson interference, Müller’s ratchet, and background selection^40–41^. A reduced *N*_e_ is predicted to allow accumulation of repetitive elements under weak natural selection^42^.

The establishment of large non-recombining regions may involve expansion of smaller regions, and it has been suggested that this may reflect selection maintaining linkage between initially small sex-determining regions and sexually antagonistic (SA) polymorphisms in the PAR of the chromosome carrying them^43^. Recently, alternatives to the SA) polymorphism hypothesis have been modelled, including a model involving the simultaneous loss of Y gene functionality or expression and evolution of dosage compensation^44^. However, recombination suppression has also been observed in evolutionarily young sex chromosomes with limited sequence divergence^45^, suggesting that recombination suppression may be considered independently from genomic degeneration. Mammalian Y chromosomes, with a shared common ancestor, share several well-characterized features: their Y chromosomes are highly repetitive, and contain very few genes, with a low gene density, and they mostly AT-rich, although certain large palindromic ampliconic regions are GC-rich^46–49^. Although some features such as TE enrichment appear common to plant MSY regions that evolved independently in different lineages, these regions often have high gene densities, though formal comparative analyses are lacking.

Recent advances in artificial intelligence (AI) have produced major breakthroughs, notably in large language models (LLM). Although they were initially developed for natural language processing, this modeling framework has been extended to biological contexts. Protein language models (pLM), represented by the AlphaFold series^50–51^, have demonstrated that learning about evolutionary constraints from amino acid sequences enables accurate three-dimensional structure prediction. More recently, genome language (or DNA foundation) models (gLM), which allow comprehensive learning about genomic sequence contexts and can embed them into high-dimensional latent vector representations, have been developed. These gLMs not only treat genomes as combinations of nucleotides, but also project evolutionarily conserved patterns and contextual information into “latent embeddings” exceeding a thousand dimensions. Comparing and interpreting these embeddings may allow novel “alignment-free” analyses of evolutionary constraints that overcome limitations in alignment-based approaches. The latest gLMs handling large-scale genomic contexts, the Evo series, has been trained on genomic sequences including all domains of life^52–53^. In this study, as well as Evo2^53^, we employed PlantCAD2^54–55^, a plant-specific genome language model trained on angiosperm genomes, which has demonstrated better performance than the Evo series in analyses of evolutionary constraints and interpretation of regulatory functions in non-coding regions^55^. We analyzed the genomes of *Silene* and *Humulus*, two genera with independently evolved heteromorphic Y chromosomes.

### Two interpretations of gLM-derived embeddings

As shown in Figure 1, we collected genome-wide 2-kb fragments from *Silene* and *Humulus*, which respectively belong in the euasterid and eurosid clades, which diverged more than 100 million years ago^37–39^. We converted the data into 1,536- and 4,096-dimensional embeddings using PlantCAD2 (L-size) and Evo2 (evo2_7b), respectively, and designed downstream analyses based on chromosome-level representations obtained by average pooling of these embeddings (as detailed in the Methods). Following a previous study^56^, to minimize the influence of selective constraints specific to particular gene functions, we extracted embeddings from all 2-kb regions upstream of gene transcription start sites, treating each upstream region as one sample. We conducted two complementary interpretations of the resultant embeddings. The first approach is a hypothesis-agnostic analysis, in which angular distances between vectors pooled at the chromosome level were computed across all samples, transformed into manifold distances, and analyzed by hierarchical clustering. The second approach is a PCA-based analysis aiming to extract composite Y-specific feature dimensions, followed by characterization of associated biological properties.

**Figure 1.**
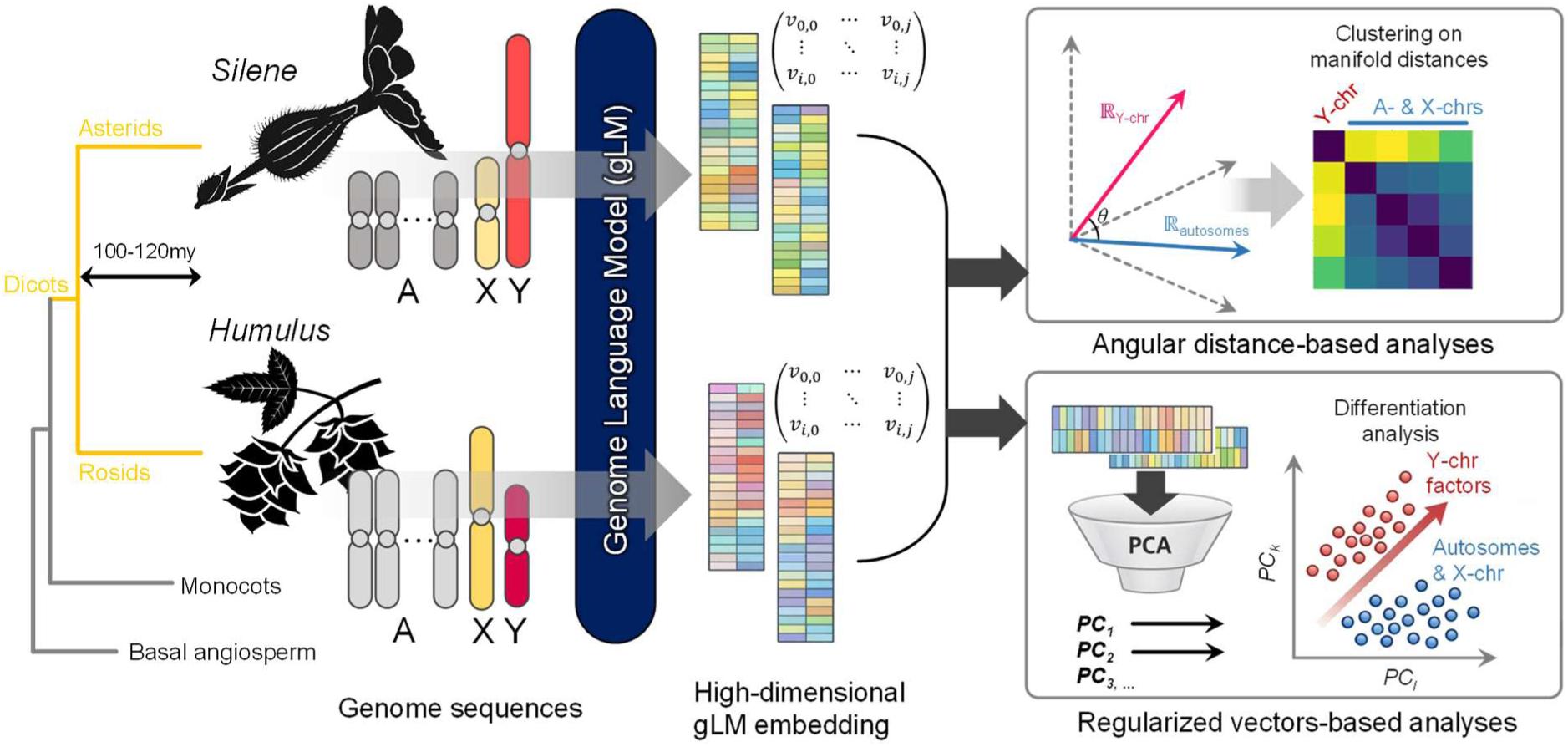
Analytical workflow for plant sex chromosomes using genome language models. In the genera *Silene* and *Humulus*, which diverged approx. 100-120 mya and independently evolved heteromorphic sex chromosomes, we investigated the convergently accumulated evolutionary constraints on the Y chromosome. Sequence features were extracted in a high-dimensional, alignment-free manner using genome language models. We applied two analytical approaches; (1) a hypothesis-agnostic method based on vector similarity using angular distance, and (2) a regularized vector-based method using principal component analysis (PCA) to detect confounding Y-specific vector components. For each approach, we further explored the sequence patterns contributing to the Y-specific vector components identified.

### Features of Y chromosomes revealed by approach 1, angular distance-based analysis

The 2-kb regions upstream of gene transcription start sites from *Humulus lupulus*, *Humulus japonicus*, *Silene latifolia*, and *Silene dioica* tended to yield species-specific clusters for all chromosomes except the MSY (Figure 2a for PlantCAD2, Supplementary Figure 1 for Evo2). Within each of these distantly related genera (though not between them), homologous chromosomes (carrying orthologous gene pairs) are shared between the two species, and embeddings were derived from upstream regions of orthologous genes. Therefore, if sequence homology were the dominant signal, homologous chromosomes within each genus should cluster together. However, this did not occur, indicating that the analysis captures lineage-specific effects as its primary features, rather than simple sequence homology. Notably, with PlantCAD2, in both *Silene* and *Humulus*, the MSYs were consistently positioned as isolated clusters or outgroups (Figure 2a), suggesting (unsurprisingly) that they have distinctive features. Because the species used to train PlantCAD2 do not include any with heteromorphic sex chromosomes, we tested whether this limitation could lead to an inadequate representation of Y chromosome features (i.e., providing only sparse information and consequently placing them as an outgroup). To examine this possibility, we performed the same analysis including mouse Y chromosome. The mouse Y chromosome was placed as a clearly distinct outgroup, separate from the Y chromosomes of *Silene* and *Humulus* (Supplementary Figure 2). This suggests that PlantCAD2 would properly capture features that are specific to the Y chromosomes of *Silene* and *Humulus*. On the other hand, when Evo2 was used, the Y chromosomes of *Silene* were positioned as a distinct outgroup, whereas the Y chromosomes of *Humulus*, although forming a cluster separate from autosomes, were distinguished from those of *Silene* (Supplementary Figure 1). This is likely because the training genome dataset of Evo2 includes *Silene latifolia* and *Silene dioica*, but does not include species from the genus *Humulus*, thereby allowing features characteristic of *Silene* Y chromosomes to be more prominently represented.

**Figure 2.**
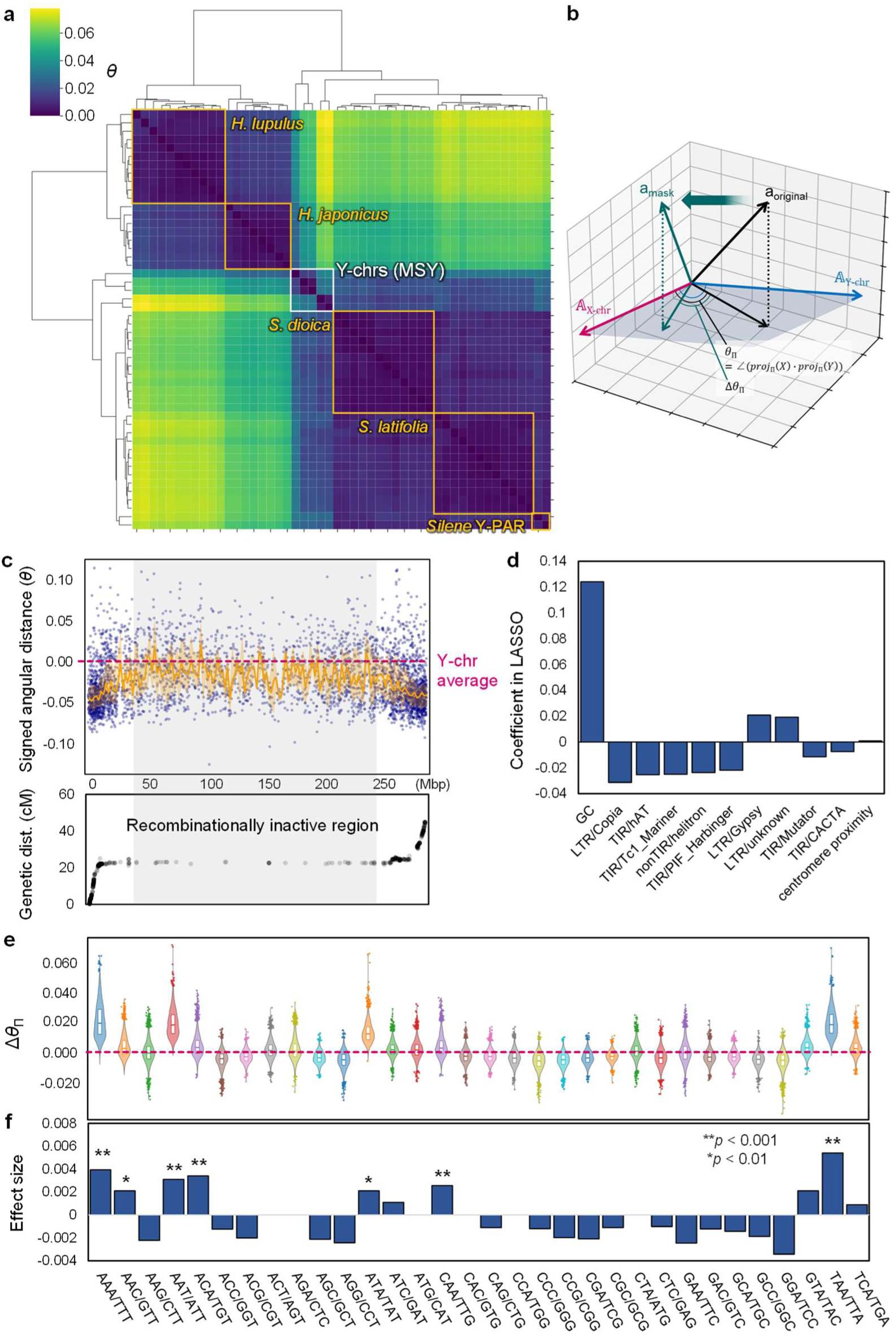
Angular distance-based detection of Y chromosome-specific properties. **a**, Hierarchical clustering based on angular distance transformed into a manifold distance. For autosomes and the X chromosome, clear species-specific clusters are formed (outlined in orange and annotated with species names), and their relationships are consistent with the phylogenetic relationships. In contrast, the MSY region of the Y chromosome forms an independent subclade that includes members of both the *Silene* and *Humulus* genera (outlined in white and annotated as Y-chr (MSY)). **b,** Conceptual illustration of the angular distance (*θ*_Π_) under projection onto the subspace defined by the X and Y chromosome vectors, and the change in angular distance (Δ*θ*_Π_) upon masking within this X-Y subspace. **c,** Trajectory of the signed angular distance in the X-Y subspace (*θ*_Π_) along the X chromosome of *Silene latifolia* (top), and a Marey map showing the relationship between genetic and physical distance (bottom). Pericentromeric recombinationally inactive regions (highlighted in gray) exhibit values closer to those of the Y chromosome. **d,** Comparison of coefficients from LASSO regression using *θ*_Π_ as the response variable in *S. latifolia*. Representative genomic properties, including GC content, density of various transposable elements, and proximity to the centromere, were included as explanatory variables. **e-f,** Distribution of Δ*θ*_Π_ upon masking each 3-mer (shown as violin plots with median and first/third quartiles indicated by box plots) (**e**), and the effect size of each 3-mer masking estimated by a Bayesian mixture model, with asterisks indicating posterior probabilities of *p* < 0.01 (*) or < 0.001 (**) (**f**).

To examine whether these Y-specific features arise from localized sequence motifs, we performed clustering after emphasizing local signals within individual vectors using GemPooling (P = 1, 2, 4, and 6), where P = 1 corresponds to standard average pooling, and increasing P progressively emphasizes localized signals, approaching max-pooling-like behavior. However, greater local signal enhancement (or higher P), caused increasingly uncertain overall cluster structure (Supplementary Fig. 3). Y specificity thus does not reflect specific motif signals but some kind of wider patterns.

To further interpret the Y-specific features, we focused on *Silene latifolia*, the species whose Y is best characterized. We defined a subspace of the pooled X and Y chromosome vectors and projected individual vectors on this subspace to detect the projected angular distance (*θ*_Π_, not to be confused with the commonly used notation for nucleotide diversity), which simulates distances to the Y chromosome in the plane directly connecting the X and Y vectors (Figure 2b). Along the X chromosome, *θ*_Π_ values correlated closely with recombination rates (Figure 2c for PlantCAD2, Supplementary Fig. 4 for Evo2). Specifically, the oldest recombinationally inactive region in XY males, which occupies a physically large central part of the X chromosome^31–33^, exhibited *θ*_Π_ values closer to those of the Y chromosome than the rest of the chromosome. When vectors from recombinationally active and inactive regions of the X chromosome were projected onto the orthogonal complement of their subspace, which should remove the effects of recombination suppression, the distance between X and MSY was indeed substantially reduced (Supplementary Figure 5). Nevertheless, the MSY still formed a distinct cluster, suggesting that MSY-specific features are not explainable solely by the sequence pattern derived from recombination suppression.

In *Silene latifolia*, gradual expansion of recombinationally inactive regions is reflected in so-called “evolutionary strata” of differing Y-X sequence divergence in different X chromosome regions, similar to the strata originally discovered in the human XY pair^57^. At least two strata (S0 and S1) are inferred in *S. latifolia*, using synonymous site divergence between coding sequences of single-copy genes with copies on both the X and Y chromosomes (termed gametolog pairs, which can be identified, despite the Y chromosome being highly rearranged); the older S0 and younger S1 strata were defined as having *dS_XY_* values above and below 11%, respectively (with ranges from 0.11 to 0.21 versus 0.03 to 0.11, according to Akagi et al. 2025^32^). We analysed *θ*_Π_ separately for these strata, including clustering based on their *dS_XY_* values (Supplementary Fig. 6). The older recombination-suppressed stratum, S0, exhibited lower *θ*_Π_ values and was closer to the autosomes than to the S1 region. At first sight, this appears unexpected, but it might reflect different ancestral states predating X-Y divergence. To further investigate this possibility, we analysed genome-wide *θ*_Π_ in *S. vulgaris*, which diverged from *S. latifolia* before its sex chromosomes evolved^32–33^, and can therefore serve as an outgroup likely to have retained the ancestral states of genome regions. The *S. vulgaris* region corresponding to half of *S. latifolia*’s S1 is made up of recombinationally inactive regions of chromosomes 2 and 3, while its other half corresponds to a large recombinationally active part of the right arm of the *S. vulgaris* chromosome 4^32–33^. On the other hand, the *S. vulgaris* region corresponding to *S. latifolia*’s S0 is made up of recombinationally further active part of the right arm of the *S. vulgaris* chromosome 4^32–33^. Consistent with their genetic properties, *θ*_Π_ values for the recombinationally inactive chromosome 2 and 3 regions in *S. vulgaris* (corresponding the half of S1) were substantially elevated compare with the recombinationally active part of the right arm of *S. vulgaris* chromosome 4 (corresponding the other half of S1 and the whole S0) (Supplementary Fig. 7). Together, these results suggest that the observed differences in *θ*_Π_ values between S0 and S1 might not primarily reflect the timing of recombination suppression of the Y chromosome. Rather, it may reflect ancestral chromosomal states pre-dating the X-Y recombination suppression that created the S0 region, while the region that became the younger S1 stratum probably already recombined rarely and had a high repeat density^32^.

To identify specific sequence contexts contributing to Y chromosome specificity, we first considered features known from mammalian Y chromosomes and performed LASSO regression with *θ*_Π_ as the response variable and GC content, coverage by various transposable elements, and proximity to centromeres as explanatory variables. GC content showed the strongest positive contribution toward the Y direction, while positive contributions were also observed for coverage by *Gypsy* and *unknown*-class LTR-type transposable elements (Figure 2d for PlantCAD2, Supplementary Fig. 8 for Evo2). *Gypsy* and *unknown* elements are known to have accumulated specifically and on this Y chromosome^32^, consistent with our findings. No significant multicollinearity was detected among the different explanatory variables (variance inflation factor < 1.4). There thus appears to be a strong independent effect of GC content.

Next, we performed an occlusion sensitivity analysis using random 3-mer masking for each sequence, and quantified changes in *θ*_Π_ (Δ*θ*_Π_ in Figure 2b). Simple masking effects were strongest for AT-rich 3-mers (Figure 2e). However, after correction with GC content effects and category effects of the original sequence groups as covariates in a Bayesian mixture model, in addition to AT-rich motifs, 3-mers composed of purine-pyrimidine combinations (CAA/TTG, ACA/TGT, AAC/GTT) showed consistently high contribution probabilities (posterior prob. > 0.99; Figure 2f). This suggests that these motifs are significantly depleted on the MSY. Genome-wide distributions of GC content and CAA motifs revealed that (old) recombination-suppressed regions derived from pericentromeric regions tended to exhibit high GC content (Supplementary Fig. 9, *p* = 7.9e^-34^ for recombinationally active vs inactive regions, in Mann-Whitney U tests) and faintly low CAA abundance (Supplementary Fig. 10, *p* = 4.5e^-8^ for recombinationally active vs inactive regions, in Mann-Whitney U tests), indicating a potential association between these features. Importantly, the MSY showed further exaggerated features than pericentromeric non-recombining regions of any other chromosome (higher GC content and lower CAA abundance were significant by Mann-Whitney U tests, with *p* < 1e^-64^).

### Features of Y chromosomes revealed by approach 2, Y-chromosome-specific regularized vectors

The angular distance-based analysis described above was hypothesis-agnostic and applied to all vector components. In the second set of analyses, we instead aimed to intentionally extract Y-chromosome-specific composite principal components and to interpret genomic properties along these axes. We performed principal component analysis (PCA) on chromosome-level 1,536- and 4,096-dimensional feature vectors obtained by average pooling with PlantCAD2 and Evo2, respectively, and identified principal component axes with statistically significant differences between the MSY of *Silene* and *Humulus* and their X-chromosomes or autosomes (for PlantCAD2, PC3, *p* = 0.0016 and PC1, *p* = 0.085, respectively; Fig. 3a, for Evo2, PC2, *p* = 8.5e^-6^ and PC1, *p* = 0.0061, respectively; Supplementary Fig. 11, in Mann-Whitney U tests). In the comparison between PlantCAD2 and Evo2, the separation between the Y-chromosomes and the autosomes along the principal component axis showing Y-chromosome specificity was clearer in PlantCAD2 (see Fig. 3a and Supplementary Fig. 11); therefore, we mainly examined the features captured by PlantCAD2.

**Figure 3.**
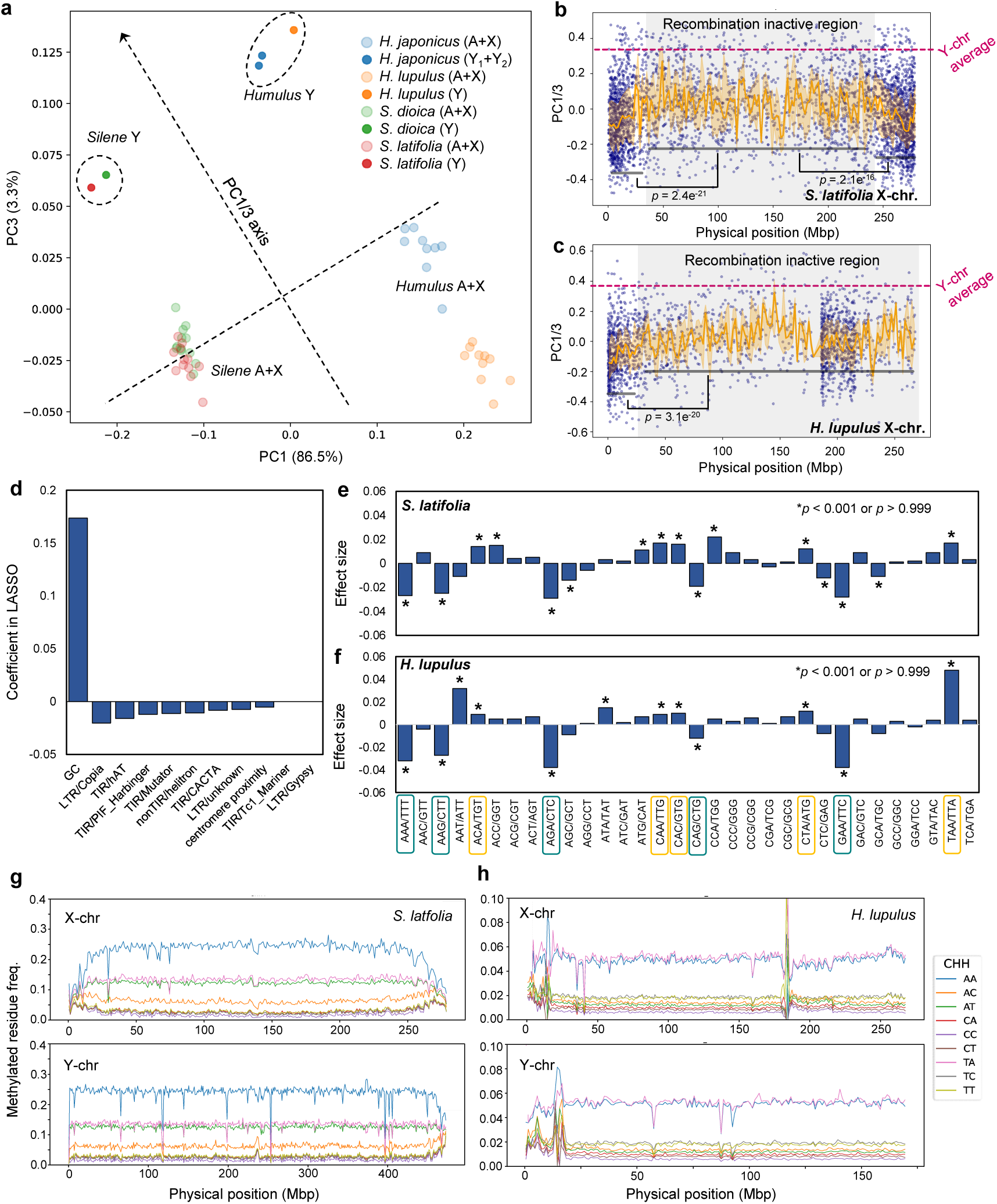
Regularized vector-based detection of Y chromosome-specific properties. **a**, Distribution of autosomes and X chromosomes (semi-transparent plots) and Y chromosomes (thick plots) for the genera *Silene* (*S. latifolia* and *S. dioica*) and *Humulus* (*H. lupulus* and *H. japonicus*) in the PC1/PC3 space. PC1 primarily contributes to species separation; however, along the combined PC1/PC3 axes, both genera show concordance in the direction and norm (or distance) of the vectors separating autosomes/X chromosomes from Y chromosomes. **b-c,** Distribution of PC1/PC3 values along the X chromosome in *S. latifolia* (**b**) and *H. lupulus* (**c**). Recombinationally inactive regions shift toward Y chromosome-like values and show significant differences compared to recombinationally active regions (p < 2.1 x 10^-16^, in Student’s *t*-test). **d,** LASSO regression using PC1/PC3 values across the whole genomes of *S. latifolia* and *H. lupulus* as the response variables. As in Figure 2d, representative genomic properties (GC content, transposable element density, and proximity to the centromere) were used as the explanatory variables. **e-f,** Comparison of effect sizes on PC1/PC3 values from 3-mer masking based on a Bayesian mixture model in *S. latifolia* (**e**) and *H. lupulus* (**f**). Asterisks indicate significant posterior probabilities (*p* < 0.001 or > 0.999). For sequence patterns showing significant posterior probabilities in both genera, those with positive effect sizes (i.e., masking brings values closer to Y chromosome properties) are outlined in orange, whereas those with negative effect sizes (i.e., masking moves values away from Y chromosome properties) are outlined in dark cyan. **g-h,** Patterns of DNA methylation bias in CHH contexts on the X and Y chromosomes of *S. latifolia* (**g**) and *H. lupulus* (**h**). In both species, CAA methylation is consistently predominant.

In both *Silene* and *Humulus*, PC1 mainly reflected inter-species divergence, whereas PC3 consistently separated the Y chromosome from the autosomes and the X, in both genera. Along the mixed PC3/PC1 axis (denoted PC1/3 axis, and indicated by dashed lines in Fig. 3a), vectors from autosomes and X chromosomes to Y chromosomes exhibited consistent directions and nearly identical norms (or distance) in both genera. In contrast, when we explored principal components that showed significant differences between the X chromosome and the others, PC12 and PC16 were identified as important (*p* = 0.018 and 0.011, respectively, in Mann-Whitney U tests); however, along these axes, the X chromosome did not form a distinct cluster relative to the other chromosomes (Supplementary Fig. 12).

The recombinationally inactive regions of the X and autosomes exhibited significantly elevated PC1/3 values, approaching those of the Y chromosome (Fig. 3b-c for X chromosomes of *Silene* and *Humulus*; Supplementary Fig. 13 for genome-wide statistic tests, *p* = 1.4e^-153^ in Student’s *t*-test for recombinationally active vs. inactive regions in *Silene latifolia*). In *Silene latifolia*, the pericentromeric region of the X chromosome (with a long history of old recombinational inactivity), showed high PC1/3 values, strongly shifted toward those of the Y chromosome (Fig. 3b). This observation is consistent with the angular distance results described above, and supports the view that Y-chromosome properties basically reflect long-term recombination suppression. Note that, in Evo2, PC1/2 values exhibited consistent trajectory (Supplementary Fig. 14). To further investigate this pattern, we compared PC1/3 values in PlantCAD2 between strata with different times since recombination suppression: S0 and S1 in *S. latifolia*, and S0-1 and S2 in *Humulus japonicus*. In *H. japonicus*, the older strata (S0-1; *dS* = 0.21-0.32, according to Akagi et al. 2025^32^) exhibited values exceeding the Y-chromosome average, whereas the much younger stratum S2 (*dS* = 0.03-0.08) showed only a modest increase (Supplementary Fig. 15). In contrast, in *S. latifolia*, PC1/3 values were higher in the younger S1 than in the older S0 stratum, consistent with the suggestion above that they reflect ancestral recombinational states rather than the time since complete cessation of X-Y recombination stopped, initiating X-Y divergence.

To identify contributing attributes, we performed LASSO regression using GC content, TE classes, and proximity to centromeres as explanatory variables across all genomic components in *Silene* and *Humulus*. Consistent with the angular distance analysis, GC content showed the strongest positive contribution (Fig. 3d). We next conducted 3-mer random masking sensitivity analyses separately for *Silene* and *Humulus* to quantify changes in PC1/3 values (ΔPC1/3). Using a Bayesian mixture model that takes account of GC content and the original sequence category as covariates associated with masking, we identified motifs with strong positive contributions (posterior prob. > 0.999) shared by both genera, including ACA/TGT, CAA/TTG, CAC/GTG, CTA/TAG, and TAA, as well as motifs with strong negative contributions, including AAA/TTT, AAG/CTT, AGA/TCT, CAG/CTG, and GAA/TTC (Fig. 3e-f). These results, are mostly consistent with the results of the angular distance analysis in suggesting depletion of A-C or T-G pairs on the Y chromosome, accompanied by an accumulation of purine-purine and pyrimidine-pyrimidine combinations.

### Interpretation of plant-specific Y-chromosome properties

Here, we integrate the properties shared between the Y chromosomes of the distantly related genera *Silene* and *Humulus*, as inferred from genome language model embeddings, with existing understanding of genome and sex chromosome evolution. As outlined in the Introduction section, heteromorphic Y chromosomes are expected to have undergone recombination suppression events, leading to Hill-Robertson interference that weakens natural selection, resulting in Y chromosome degeneration, with genes having lower functionality than their X gametologs, or being deleted. The observed depletion of purine-pyrimidine combination motifs such as C-A (or G-T), accompanied with the retention of purine-purine and pyrimidine-pyrimidine combinations such as C-T (or G-A), may therefore reflect background patterns associated with neutral epigenomic modifications. In plants, cytosine methylation is known to exhibit sequence-context biases, particularly within the highly abundant CHH context among the three methylation contexts (CG, CHG, and CHH)^58^. Consistent with this, both *S. latifolia* and *H. lupulus* showed the highest DNA methylation frequencies in CAA sequences, followed by CTA (and 3-mers starting from “CA” in *Silene*) (Fig. 3g-h for the X and Y chromosomes). Importantly, these sequence patterns mostly correspond to those whose masking contributes to the Y-specific property (see Fig. 2f, Fig. 3e-f). Methylated cytosines are prone to deamination, resulting in preferential substitution to thymine, a major driver of *de novo* SNP in plants^59^. In completely sex-linked regions of sex chromosomes, which in *S. latifolia* are typically TE-rich and highly methylated^32–34^, the effects of cytosine deamination are likely to be pronounced, and may persist unless opposed by strong natural selection, as proposed in Fig. 4. Hence, biased cytosine methylation may explain the significant depletion of CAA (or TTG) (and CTA or CA-related) contexts. In vertebrate genomes, methylated CpGs undergo spontaneous deamination to thymine, leading to elevated rates of C to T and G to A substitutions that reduce GC content over evolutionary time^60–61^. Such mutational biases are expected to be particularly prominent in non-recombining and weakly selected genomic regions such as the MSY.

**Figure 4.**
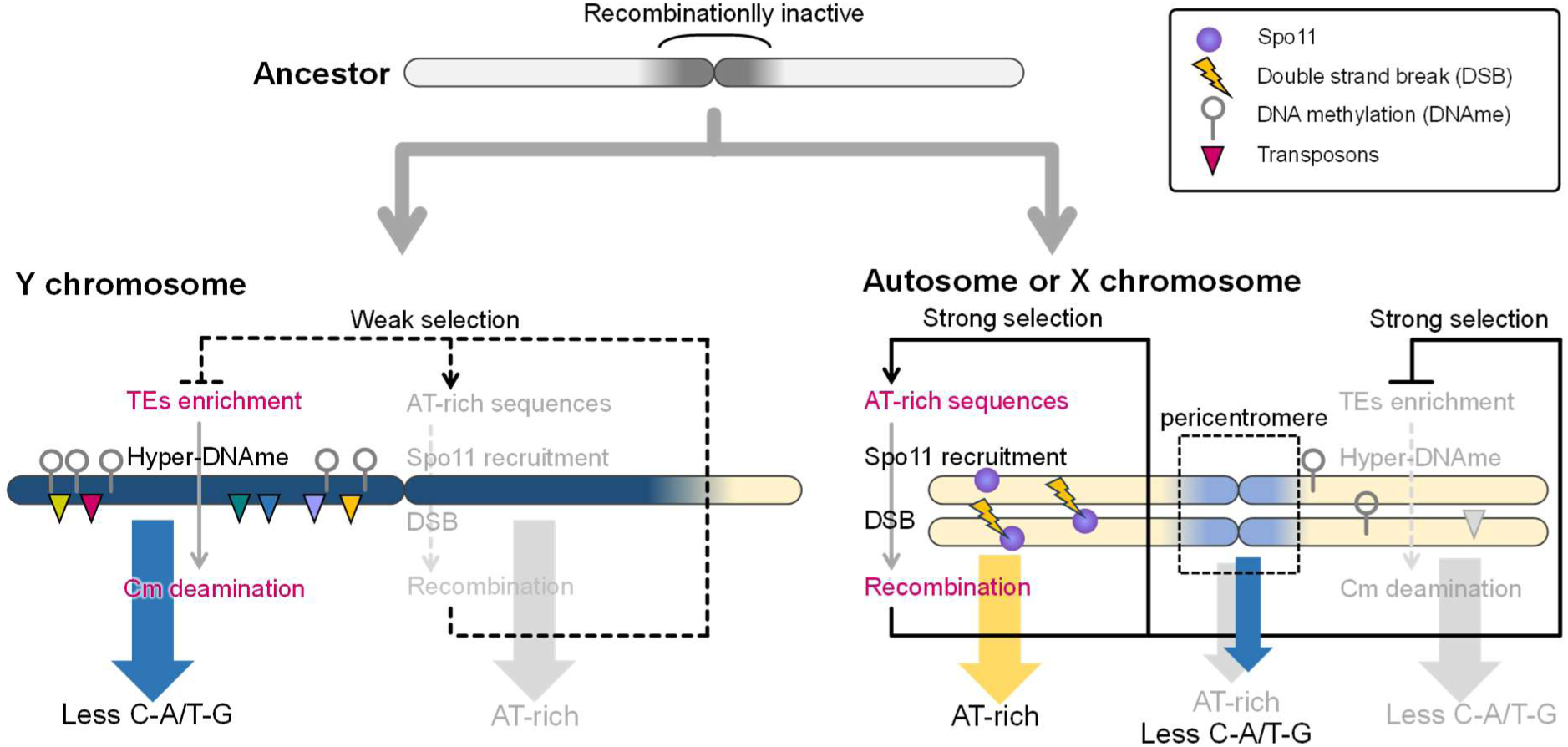
A model for the establishment of Y chromosome-specific context. Even in ancestral chromosomes prior to the emergence of sex chromosomes, recombination suppression is already advanced in pericentromeric regions. Following the divergence into autosomes (or X chromosomes) and Y chromosomes, AT-rich regions in autosomes undergo recognition by Spo11, promoting double-strand breaks (DSB) and thereby activating recombination. This recombination activity enables natural selection to act efficiently, resulting in a positive feedback loop that maintains AT-rich sequences. Simultaneously, appropriate removal of TEs occurs, reducing the extent of DNA hypermethylation. Consequently, the frequency of deamination in C-A/T-G sequence contexts, where DNA methylation is more likely to occur, is reduced, and loss of these sequence contexts is prevented. In contrast, on the Y chromosome, recombination suppression arises, partly driven by a substantial reduction in effective population size (*N*_e_). As a result, the positive feedback observed in autosomes (AT-richness → recombination activation → natural selection) does not operate. Instead, GC content increases relative to autosomes, TE removal becomes inefficient, and widespread DNA hypermethylation accumulates. As a consequence, loss of C-A/T-G contexts is observed, reflecting sequence contexts most strongly influenced by this hypermethylation.

However, although this framework may be consistent with the globally low GC content of mammalian Y chromosomes, it contradicts the elevated GC content observed here in plant Y chromosomes and in autosomal regions that recombine rarely. In humans, GC-biased gene conversion within large palindromic regions of the Y chromosome opposes GC loss^62^. However, neither *Silene* nor *Humulus* Y chromosomes have large palindromic structures, and this process is not expected to occur in these or other rarely recombining regions in their genomes. Therefore additional causal mechanisms must be considered.

For this issue, the sequence characteristics of plant-specific recombinationally active regions are likely to be important. AT-rich sequences increase DNA curvature and enhance chromatin accessibility^63^. In addition, AT-rich sequences are known to recruit Spo11, which induces double-strand breaks (DSBs) that promote recombination during meiosis^64–66^. As a result, recombinationally active regions are suggested to be enriched with AT-rich motifs. In studies of recombination hotspots in plants, although the enriched sequence contexts vary somewhat independently among species, many contain AT-rich motifs^67–68^. In other words, in autosomes where recombination is maintained, recombination frequency is preserved with AT-rich sequences as initiation sites. This allows avoidance of Hill-Robertson interference and enables proper natural selection to function, thereby maintaining AT-rich sequences that serve as recombination initiation sites (as a form of positive feedback), while also suppressing the accumulation of deleterious TEs (Fig. 4). In contrast, on the Y chromosome, where weakened selection is assumed, AT-rich sequences are not maintained and recombinational activity is not preserved. This may act as a negative feedback loop, further weakening selection and leading to rapid suppression of recombination. Furthermore, under weak selection, TEs are not efficiently removed, and the increase in methylated cytosine may further promote the loss of C-A (T-G) context as a background pattern (Fig. 4).

## METHODS

### Genome sequences and extraction of gene promoter regions

For the heterogametic male (XY-type) dioecious species *Silene latifolia* and *S. dioica*, and their closely related gynodioecious species *S. vulgaris*, we obtained the whole-genome sequences of male individuals (SLGerPM1 and SdJapan_07M for *S. latifolia* and *S. dioica*, respectively) and a hermaphrodite individual (SvPH1 for *S. vulgaris*)^32^ (https://genome.kazusa.or.jp/species/?kgp_id=t37657.G002 for *S. latifolia*, https://genome.kazusa.or.jp/species/?kgp_id=t39879.G001 for *S. dioica*, and https://genome.kazusa.or.jp/species/?kgp_id=t42043.G001 for *S. vulgaris*). Similarly, for the distantly related heterogametic male (XY-type) dioecious species *Humulus lupulus* (or common hop) and *Humulus japonicus* (order Rosales, eurosids), we obtained whole-genome sequences of male individuals (10_12 and Seta-line, respectively)^34^ (https://genome.kazusa.or.jp/species/?kgp_id=t3486.K002 for *H. lupulus* and https://genome.kazusa.or.jp/species/?kgp_id=t3485.K001 for *H. japonicus*). For all species, 2-kbp regions upstream of all annotated genes (5′ promoter regions) were extracted in FASTA format by using samtools^69^.

### Embedding generation by genome language models

Embeddings were generated from the FASTA files described above using PlantCAD2 (large)^55^ (https://huggingface.co/kuleshov-group/PlantCAD2-Large-l48-d1536 for model checkpoint) and evo2_7b^53^ (https://github.com/arcinstitute/evo2). The output shapes (sequence length, embedding dimension) were (2,000, 1,536) for PlantCAD2 and (2,000, 4,096) for Evo2. For evo2_7b, embeddings were extracted from the layer blocks.28.mlp.l3. The GPU environments used for embedding generation were NVIDIA RTX A5000 x 1 for PlantCAD2 and NVIDIA RTX A6000 Ada x 1 for Evo2. Promoter regions shorter than 2 kbp and low-confidence regions (those containing more than 30% ambiguous bases, N) were excluded from the analysis.

For each chromosome in each plant species, embeddings from all records were averaged along the sequence dimension (by average pooling) to to generate a single representative vectors 𝐯 ∈ ℝ^M^((1, 1536) for PlantCAD2 and (1, 4096) for Evo2). These vectors provide a compact representation of sequence composition and higher-order contextual features learned by the model. As described in the main text, we also conducted preliminary analyses using GeM pooling to modulate the weighting of local motif signals.

### Detection of angular distance based on highly dimensional vectors

To quantify similarity between sequence embeddings, we computed angular distances between embedding vectors. For two vectors 𝐯_i_and 𝐯_2_, cosine similarity was first calculated as

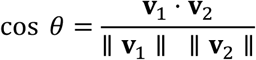

where denotes the inner product and ∥∙⋅∥∙ denotes the Euclidean norm. Cosine similarity was computed using the NumPy library (np.dot, np.linalg.norm). The angular distance 𝜃 was then defined as

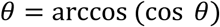

which ranges from 0 to 𝜋, corresponding to identical and opposite directions in the embedding space, respectively. Angular distance was used as a scale-invariant measure of similarity, capturing differences in vector direction rather than magnitude. This is particularly suitable for high-dimensional embedding spaces, where biologically meaningful information is often encoded in directional structure.

Given a target vector 𝐭 and a reference vector 𝐫, we defined a two-dimensional subspace spanned by these vectors (in this manuscript we applied them to the X and Y chromosomes). An orthonormal basis 𝐔 ∈ ℝ^M×2^ for this subspace was constructed using the Gram-Schmidt process (implemented in NumPy), ensuring that the basis vectors are mutually orthogonal and normalized. All embedding vectors 𝐯 were then projected onto this subspace as

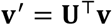

yielding a two-dimensional representation that captures variation along the target-reference axis. Angular distances were subsequently computed within this projected subspace. Each projected vector 𝐯^’^ was re-normalized to unit length, and the angular distance between two vectors 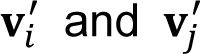 (*θ*_Π_ in the main text) was defined as

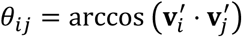

This procedure isolates the component of variation explained by the target-reference relationship while removing orthogonal contributions, enabling direct comparison of samples along the axis between X and Y chromosomes.

### Hierarchical clustering with manifold (geodesic) distance estimation from angular distances

To capture the intrinsic geometry of the embedding space, pairwise angular distances were transformed into manifold (geodesic) distances using a k-nearest neighbor (kNN) graph approach. A k-nearest neighbor graph was constructed using NetworkX. Starting from the angular distance matrix, a graph was constructed in which each sample was connected to its *k* nearest neighbors (here, *k* = 10), with edge weights defined by the angular distances. The geodesic distance between two samples was then defined as the shortest path length on this graph, computed using Dijkstra’s shortest path algorithm. This procedure approximates distances along the underlying data manifold rather than in the ambient high-dimensional space, thereby preserving nonlinear relationships among samples. The resulting distance matrix was used for hierarchical clustering. The distance matrix was converted into condensed form and subjected to agglomerative clustering using an average linkage method. Then, clustering results were visualized as a heatmap with an associated dendrogram, reflecting the manifold distance space.

### Principal component analysis (PCA) to identify Y chromosome-specific vector components

Embedding vectors derived from PlantCAD2 or Evo2 were subjected to principal component analysis (PCA) using the implementation in scikit-learn. Vectors from the target group (Y chromosome in this study) and background group (autosomes and X chromosome) were concatenated and projected onto the top 40 principal components. The resulting PCA scores for each sample, along with the explained variance ratio of each component, were recorded. To identify principal components that distinguish Y chromosome sequences from autosomal and X chromosome sequences, statistical comparisons were performed for each PC dimension. Specifically, PC scores for the target and background groups were compared using Welch’s *t*-test (two-sided, unequal variance) as implemented in SciPy. PCs showing significant differences (*p* < 0,01) were further interpreted as contributing to Y chromosome-specific variation in the embedding space.

### LASSO regression

To identify genomic features associated with embedding-derived metrics, we performed LASSO regression using GC content, densities of TEs, and proximity to the centromere as explanatory variables. The response variable was defined either as the angular distances projected onto the X-Y subspace (*θ*_Π_) or as the value along PC axes that showed significant separation of Y chromosome. All explanatory variables were standardized to zero mean and unit variance prior to model fitting using scikit-learn. Model regularization strength (α) was determined by 10-fold cross-validation using LASSO with coordinate descent (LassoCV), and the final model was refit using the optimal α. Multicollinearity among explanatory variables was preliminarily evaluated using variance inflation factors (VIF) computed with statsmodels. Regression coefficients and VIF values were used to interpret the relative contributions of genomic features associated with Y chromosome-specific properties.

### Occlusion sensitivity analysis and Bayesian mixture model for masking-specific effects

To evaluate the contribution of local sequence contexts to embedding-derived properties, we performed systematic masking of all possible 3-mer sequences (including their reverse complements) in genomic sequences. For each masked sequence, embeddings were recomputed using the same gLM as in the main analysis, and the resulting change in angular distance in the X-Y subspace (Δ*θ*_Π_) or PCs that significantly separate Y chromosomes (ΔPC1/3 for PlantCAD2 and ΔPC1/2 for Evo2) were quantified. For each masking event, we recorded the 3-mer category, the change in GC content (ΔGC), the number of masked sites (mask count), and the originating sequence. This allowed us to construct a dataset linking sequence perturbations to changes in embedding geometry.

To statistically quantify the effect of 3-mer masking while accounting for confounding variables, we employed a Bayesian mixture model implemented in PyMC. Here, we exemplified a case where the response variable was the angular distance change Δ𝜃_i_ for observation 𝑖, modeled as:

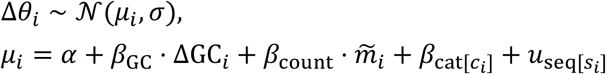

where 𝛼 is the intercept, 𝛽_GC_ and 𝛽_count_ are fixed effects for ΔGC and standardized mask count 𝑚∼_i_, respectively, 𝛽_cat_ represents category-specific effects for each 3-mer class 𝑐_i_ , and 𝑢_seq_ is a random effect accounting for sequence-specific variation 𝑠_i_ . Category and sequence effects were modeled as hierarchical priors:

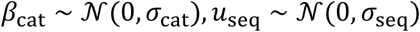

with hyperpriors 𝜎, 𝜎_cat_, 𝜎_seq_ ∼ HalfNormal(0.05), and weakly informative priors for fixed effects 𝛼, 𝛽_GC_, 𝛽_count_ ∼ 𝒩(0,0.05). Posterior distributions were estimated using Markov chain Monte Carlo (MCMC) sampling (4 chains, 2,000 tuning steps, 2,000 draws per chain). Category-specific effects were summarized by posterior means, and the posterior probabilities 𝑃(𝛽_cat_ > 0) and 𝑃(𝛽_cat_ < 0) were used to assess the direction and significance of each 3-mer effect.

### Synteny analysis

Gene-based alignments were performed between the sex chromosomes of *S. latifolia* and the whole genome of the closely related gynodioecious species *S. vulgaris*, which lacks sex chromosomes, using BLASTP (e-value < 1e^-20^, --max_target_seqs 1). The correspondence of homologous genes obtained from BLASTP, together with gene positional information from gff files, was used to generate collinearity files with MCScanX^70^ to define gene order-based synteny blocks. Syntenic blocks were subsequently visualized using SynVisio (https://synvisio.github.io/<u>#/</u>).

### Detection of DNA methylation bias in CHH contexts

DNA methylome datasets from male individuals of *S. latifolia* and *H. lupulus*, obtained in previous studies^32,34^, were processed using methylpy to generate allc files, according to the previous report^34^. From these allc files, only cytosine positions in the CHH context were extracted, and weighted DNA methylation levels were calculated for each HH dinucleotide context within 500-1000 kbp genomic bins, following the approach described previously^71^.

## ACKNOWLEDGEMENTS

We thank Prof. Deborah Charlesworth (University of Edinburgh, UK) for helpful suggestions and discussion. This work was supported by the RIKEN-TRIP initiative Field Omics to TA and GT, and in part by JSPS KAKENHI Grant Number 25K10208 to JT.

## Authors contributions

Conceptualization: TA, GT

Methodology: TA, HM, JT, GT

Investigation: TA, JT

Visualization: TA

Funding acquisition: TA, GT

Project administration: TA, GT

Supervision: TA, GT

Writing – original draft: TA

Writing – review & editing: TA, HM, JT, GT

## REFERENCES

1. Berta, P., et al. Genetic evidence equating SRY and the testis-determining factor. Nature 348, 448–450 (1990).

2. Koopman, P., et al. Male development of chromosomally female mice transgenic for Sry. Nature 351, 117–121 (1991).

3. Renner, S. S. The relative and absolute frequencies of angiosperm sexual systems: dioecy, monoecy, gynodioecy, and an updated online database. Amer J Bot 101, 1588–1596 (2014).

4. Ming, R., Bendahmane, A., Renner, S. S. Sex chromosomes in land plants. Annu Rev Plant Biol 62, 485–514 (2011).

5. Westergaard, M. The mechanism of sex determination in dioecious plants. Adv Genet 9, 217–281 (1958).

6. Charlesworth, B. & Charlesworth, D. A model for the evolution of dioecy and gynodioecy. Amer Nat 112, 975–997 (1978).

7. Charlesworth, D., Charlesworth, B., Marais, G. Steps in the evolution of heteromorphic sex chromosomes. Heredity 95, 118–128 (2005).

8. Charlesworth, D. Plant sex chromosome evolution. J Exp Bot 64, 405–420 (2013).

9. Charlesworth, D. Plant contributions to our understanding of sex chromosome evolution. New Phytol 208, 52–65 (2013).

10. Charlesworth, D. Young sex chromosomes in plants and animals. New Phytol 224, 1095–1107 (2019).

11. Barrett, S. C. & Hough, J. Sexual dimorphism in flowering plants. J Exp Bot 64, 67–82 (2013).

12. Blackburn, K. B. Sex chromosomes in plants. Nature 112, 687–688 (1923).

13. Winge, O. On sex chromosomes, sex determination and preponderance of females in some dioecious plants. CR Trav Lab Carlsberg 15, 1–26 (1923).

14. Filatov, D. A., Monéger, F., Negrutiu, I., Charlesworth, D. Low variability in a Y-linked plant gene and its implications for Y-chromosome evolution. Nature 404, 388–390 (2000).

15. Filatov, D. A. & Charlesworth, D. Substitution rates in the X-and Y-linked genes of the plants, Silene latifolia and S. dioica. Mol Biol Evol 19, 898–907 (2002).

16. Hobza, R., Lengerova, M., Svoboda, J., Kubekova, H., Kejnovsky, E., Vyskot, B. An accumulation of tandem DNA repeats on the Y chromosome in *Silene latifolia* during early stages of sex chromosome evolution. Chromosoma 115, 376–382 (2006).

17. Bergero, R., Forrest, A., Kamau, E., Charlesworth, D. Evolutionary strata on the X chromosomes of the dioecious plant *Silene latifolia*: evidence from new sex-linked genes. Genetics 175, 1945–1954 (2007).

18. Bergero, R. & Charlesworth, D. The evolution of restricted recombination in sex chromosomes. Trends Ecol Evol 24, 94–102 (2009).

19. Akagi, T., Henry, I. M., Tao, R., Comai, L. A Y-chromosome-encoded small RNA acts as a sex determinant in persimmons. Science 346, 646–650 (2014).

20. Akagi, T., et al. The persimmon genome reveals clues to the evolution of a lineage-specific sex determination system in plants. PLoS Genet 16, e1008566 (2020).

21. Horiuchi, A., et al. Ongoing rapid evolution of a post-Y region revealed by chromosome-scale genome assembly of a hexaploid monoecious persimmon (*Diospyros kaki*). Mol Biol Evol 40, msad151 (2023).

22. Harkess, A., et al. The asparagus genome sheds light on the origin and evolution of a young Y chromosome. Nature Commn 8, 1279 (2017).

23. Akagi, T., et al. Two Y-chromosome-encoded genes determine sex in kiwifruit. Nature Plants 5, 801–809 (2019).

24. Akagi, T., et al. Recurrent neo-sex chromosome evolution in kiwifruit. Nature Plants 9, 393–402 (2023).

25. Müller, N. A., et al. A single gene underlies the dynamic evolution of poplar sex determination. Nature Plants 6, 630–637 (2020).

26. Almeida, P., et al. Genome assembly of the basket willow, *Salix viminalis*, reveals earliest stages of sex chromosome expansion. BMC Biol 18, 78 (2020).

27. He, L., et al. Chromosome-scale assembly of the genome of Salix dunnii reveals a male-heterogametic sex determination system on chromosome 7. Mol Ecol Resour 21, 1966–1982 (2021).

28. Ma, X., et al. The spinach YY genome reveals sex chromosome evolution, domestication, and introgression history of the species. Genome Biol 23, 75 (2022).

29. She, H., et al. Evolution of the spinach sex-linked region within a rarely recombining pericentromeric region. Plant Physiol 193, 1263–1280 (2023).

30. Sacchi, B., et al. Phased assembly of neo-sex chromosomes reveals extensive Y degeneration and rapid genome evolution in *Rumex hastatulus*. Mol Biol Evol 41, msae074 (2024).

31. Yue, J., et al. The origin and evolution of sex chromosomes, revealed by sequencing of the Silene latifolia female genome. Current Biol 33, 2504–2514 (2023).

32. Akagi, T., et al. Rapid and dynamic evolution of a giant Y chromosome in *Silene latifolia*. Science 387, 637–643 (2025).

33. Moraga, C., et al. The *Silene latifolia* genome and its giant Y chromosome. Science 387, 630–636 (2025).

34. Akagi, T., et al. Evolution and functioning of an X-A balance sex-determining system in hops. Nature Plants 11, 1339–1352 (2025).

35. Carey, S. B., et al. ZW sex chromosome structure in *Amborella trichopoda*. Nature Plants 10, 1944–1954 (2024).

36. Renner, S. S. & Müller, N. A. Plant sex chromosomes defy evolutionary models of expanding recombination suppression and genetic degeneration. Nature Plants 7, 392–402 (2021).

37. Lebrija-Trejos, E., Wright, S. J., Hernández, A., Reich, P. B. Does relatedness matter? Phylogenetic density-dependent survival of seedlings in a tropical forest. Ecology 95, 940–951 (2014).

38. Jiao, Y., et al. A genome triplication associated with early diversification of the core eudicots. Genome Biol 13 doi.org/10.1186/gb-2012-13-1-r3 (2012).

39. Wang, H., et al. Rosid radiation and the rapid rise of angiosperm-dominated forests. Proc Natl Acad Sci USA 106, 3853–3858 (2009).

40. Charlesworth, B. The evolution of sex chromosomes. Science 251, 1030–1033 (1991).

41. Charlesworth, B. & Charlesworth, D. The degeneration of Y chromosomes. Phil Trans Royal Soc B 355, 1563–1572 (2000).

42. Charlesworth, B., Sniegowski, P., & Stephan, W. The evolutionary dynamics of repetitive DNA in eukaryotes. Nature 371, 215–220 (1994).

43. Rice, W. R. Sex chromosomes and the evolution of sexual dimorphism. Evolution 735–742 (1984).

44. Lenormand, T. & Roze, D. Y recombination arrest and degeneration in the absence of sexual dimorphism. Science 375, 663–666 (2022).

45. Bergero, R., Gardner, J., Bader, B., Yong, L., Charlesworth, D. Exaggerated heterochiasmy in a fish with sex-linked male coloration polymorphisms. Proc Natl Acad Sci USA 116, 6924–6931 (2019).

46. Skaletsky, H., et al. The male-specific region of the human Y chromosome is a mosaic of discrete sequence classes. Nature 423, 825–837 (2003).

47. Wilson, M. A. & Makova, K. D. Genomic analyses of sex chromosome evolution. Annu Rev Genom Human Genet 10, 333–354 (2009).

48. Cortez, D., et al. Origins and functional evolution of Y chromosomes across mammals. Nature 508, 488–493 (2014).

49. Hughes, J. F. & Page, D. C. The biology and evolution of mammalian Y chromosomes. Annu Rev Genet 49, 507–527 (2015).

50. Jumper, J., et al. Highly accurate protein structure prediction with AlphaFold. Nature 596, 583–589 (2021).

51. Abramson, J., et al. Accurate structure prediction of biomolecular interactions with AlphaFold 3. Nature 630, 493–500 (2024).

52. Nguyen, E., et al. Sequence modeling and design from molecular to genome scale with Evo. Science 386, eado9336 (2024).

53. Brixi, G., et al. Genome modelling and design across all domains of life with Evo 2. Nature 10.1038/s41586-026-10176-5 (2026).

54. Zhai, J., et al. Cross-species modeling of plant genomes at single-nucleotide resolution using a pretrained DNA language model. Proc Natl Acad Sci USA 122, e2421738122 (2025).

55. Zhai, J., et al. PlantCAD2: A Long-Context DNA Language Model for Cross-Species Functional Annotation in Angiosperms. *bioRxiv* 10.1101/2025.08.27.672609 (2025).

56. Pearce, M., et al. Finding the tree of life in Evo 2. Goodfire Research, https://www.goodfire.ai/research/phylogeny-manifold (2025).

57. Lahn, B. T. & Page, D. C. Four evolutionary strata on the human X chromosome. Science 286, 964–967 (1999).

58. Gouil, Q. & Baulcombe, D. C. DNA methylation signatures of the plant chromomethyltransferases. PLoS Genet 12, e1006526 (2016).

59. Ossowski, S., et al. The rate and molecular spectrum of spontaneous mutations in *Arabidopsis thaliana*. Science 327, 92–94 (2010).

60. Fryxell, K. J. & Zuckerkandl, E. Cytosine deamination plays a primary role in the evolution of mammalian isochores. Mol Biol Evol 17, 1371–1383 (2000).

61. Mugal, C. F., Arndt, P. F., Holm, L., Ellegren, H. Evolutionary consequences of DNA methylation on the GC content in vertebrate genomes. G3 5, 441–447 (2015).

62. Hallast, P., Balaresque, P., Bowden, G. R., Ballereau, S., Jobling, M. A. Recombination dynamics of a human Y-chromosomal palindrome: rapid GC-biased gene conversion, multi-kilobase conversion tracts, and rare inversions. PLoS Genet 9, e1003666 (2013).

63. Duan, C., et al. Reduced intrinsic DNA curvature leads to increased mutation rate. Genome Biol 19, 132 (2018).

64. Choi, K., et al. Arabidopsis meiotic crossover hot spots overlap with H2A.Z nucleosomes at gene promoters. Nature Genet 45, 1327–1336 (2013).

65. Choi, K., et al. Recombination rate heterogeneity within Arabidopsis disease resistance genes. PLoS Genet 12, e1006179 (2016).

66. Choi, K., et al. Nucleosomes and DNA methylation shape meiotic DSB frequency in *Arabidopsis thaliana* transposons and gene regulatory regions. Genome Res 28, 532–546 (2018).

67. Shilo, S., Melamed-Bessudo, C., Dorone, Y., Barkai, N., Levy, A. A. DNA crossover motifs associated with epigenetic modifications delineate open chromatin regions in Arabidopsis. Plant Cell 27, 2427–2436 (2015).

68. Kianian, P. M., et al. High-resolution crossover mapping reveals similarities and differences of male and female recombination in maize. Nature Commn 9, 2370 (2018).

69. Li, H. et al. The sequence alignment/map format and SAMtools. Bioinformatics 25, 2078–2079 (2009).

70. Wang, Y., et al. MCScanX: a toolkit for detection and evolutionary analysis of gene synteny and collinearity. Nucl Acid Res 40, e49 (2012).

71. Akagi, T. & Sugano, S. S. Random epigenetic inactivation of the X-chromosomal *HaMSter* gene causes sex ratio distortion in persimmon. Nature Plants 10, 1643–1651 (2024).

