## Supplementary Figures for "Genomic properties representing plant sex chromosome evolution interpreted with genome language models"

**Title**

**Supplementary Fig. 1**

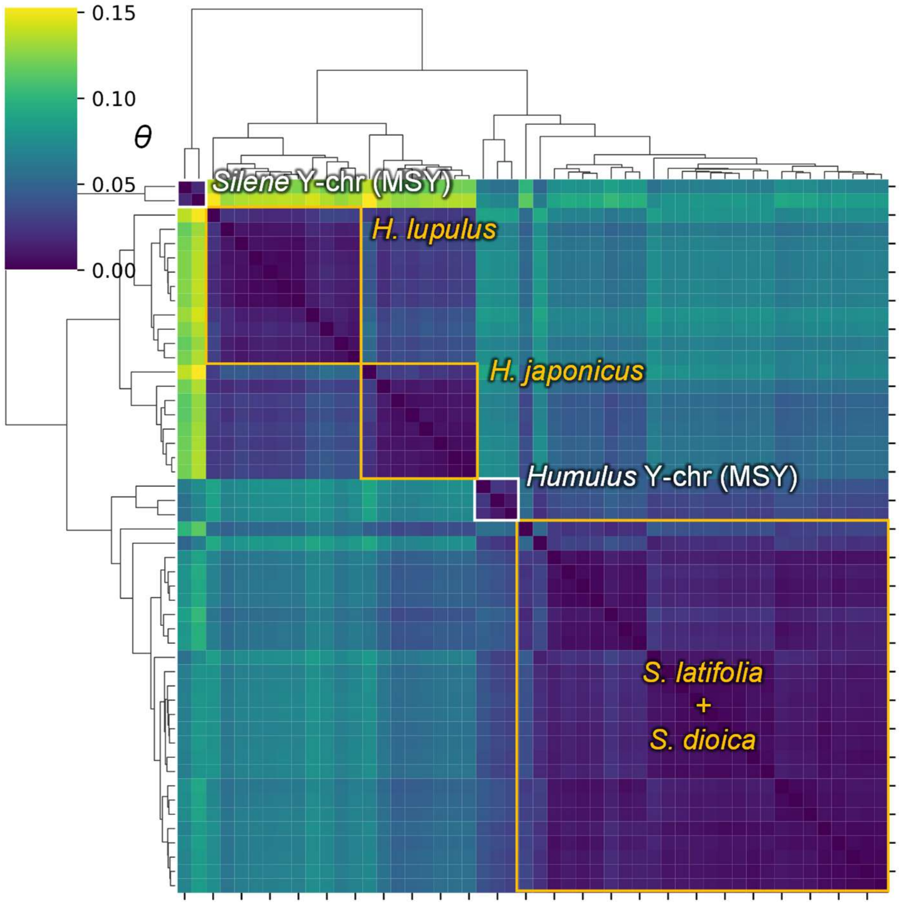


**Supplementary Figure 1 Angular distance analysis using embeddings generated by evo2**

Angular distances were calculated from embeddings generated by Evo2 (7b) and transformed into manifold distances for hierarchical clustering. As in the analysis using PlantCAD2 (Fig. 2a), species- or genus-specific clusters (outlined by orange) reflecting phylogenetic relationships were formed, and Y chromosomes (MSY) tended to be positioned as outgroups (outlined by white). However, differences between the Y chromosome components of *Silene latifolia* and *S. dioica* became more pronounced, leading to their classification into more distant outgroups. This pattern may depend on the training data of evo2: Y chromosome sequences from *S. latifolia* and *S. dioica* are included in the training dataset, whereas Y chromosome information from the genus *Humulus* has not been included.

**Supplementary Fig. 2**

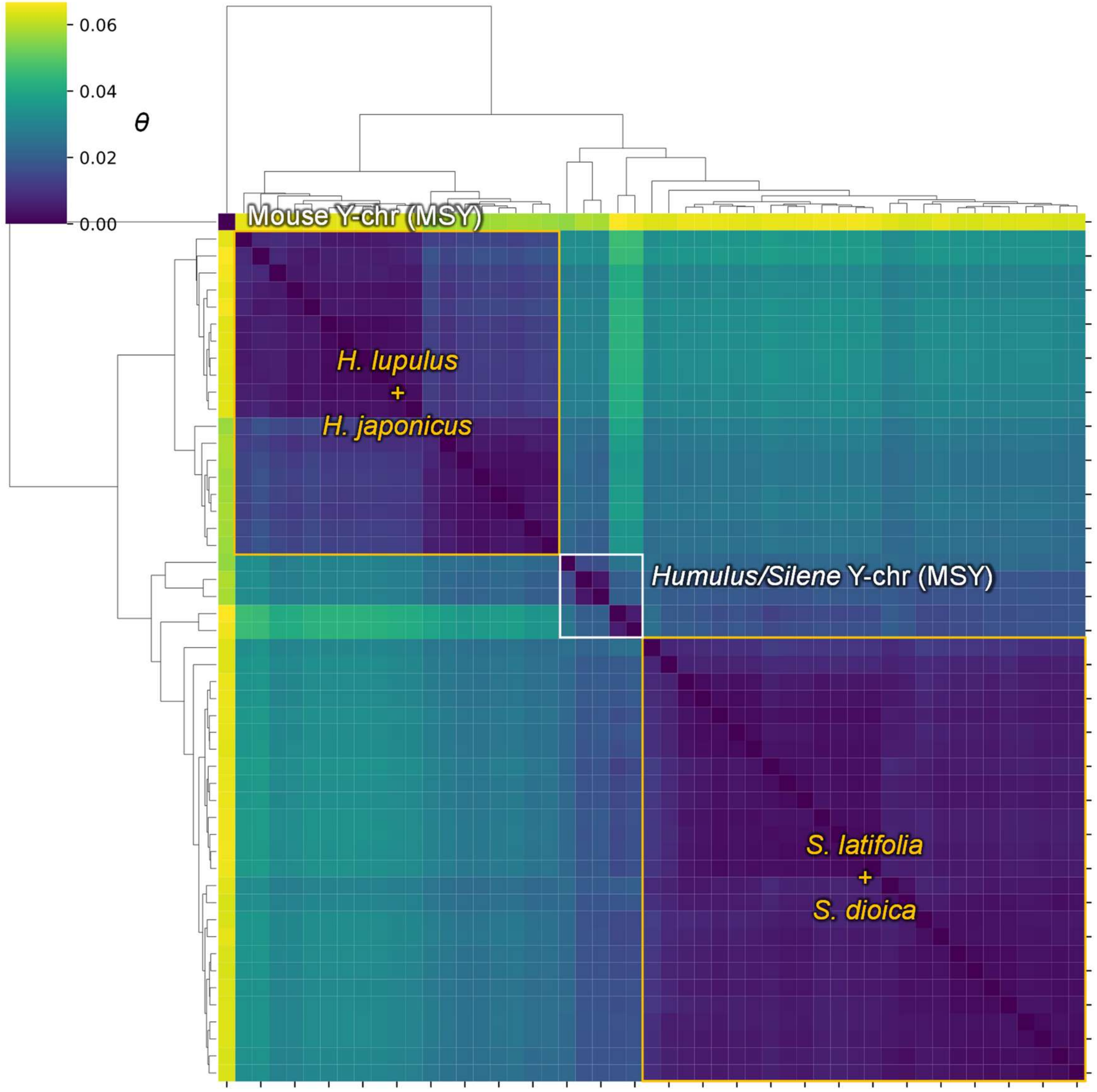


**Supplementary Figure 2 Angular distance analysis using PlantCAD2 including mouse Y chromosome data**

In the angular distance analysis shown in Figure 2a, the Y chromosomes of *Silene* and *Humulus* form a single independent cluster. To examine whether this clustering reflects a shared underlying property rather than sparsity of information due to insufficient representation of sex chromosomes in the training data, we performed an angular distance analysis including mouse Y chromosome sequences. The mouse Y chromosome clearly grouped into a cluster distinct from those of plant Y chromosomes, indicating that although plant Y chromosome sequences are not included in the training of PlantCAD2, their characteristic properties may nevertheless be captured and interpreted by the model.

**Supplementary Fig. 3**

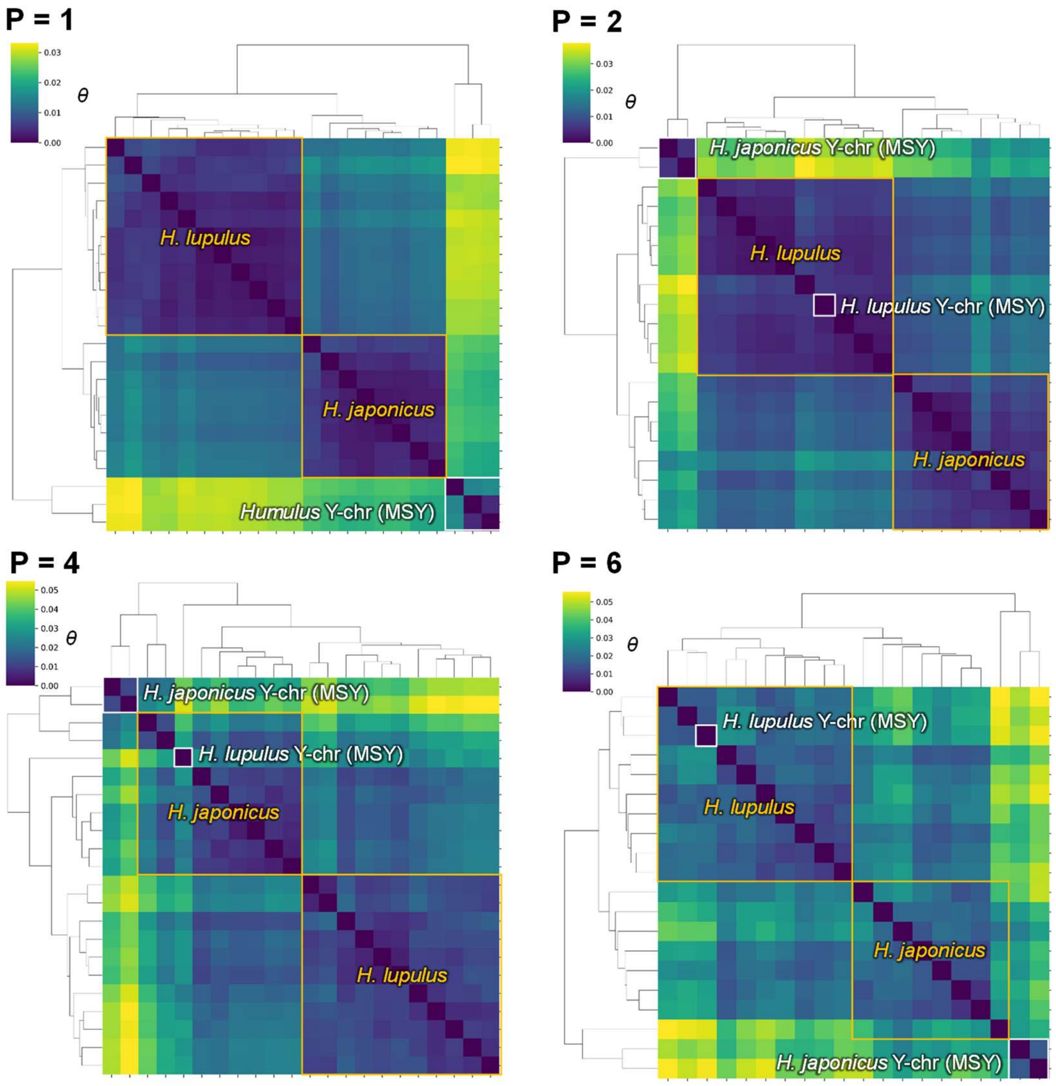


**Supplementary Figure 3 Effect of emphasizing local sequence signals by GeM pooling on angular distance clustering**

In GeM pooling, setting P = 1 corresponds to average pooling, resulting in complete averaging of residue-level embeddings. As P increases, more localized signals are increasingly emphasized. Here, we examined the relationships among angular distance clusters at P = 1,2,4, and 6 using embeddings derived from autosomes and X/Y chromosomes in the genus *Humulus*. As P increased, the distinct cluster composed solely of Y chromosomes gradually collapsed, and species-level clustering also became increasingly ambiguous. These results suggest that the sequence features defining species identity and Y chromosome specificity are not driven by discrete motif-like signals, but rather by small-scale sequence contexts distributed broadly across the genome.

**Supplementary Fig. 4**

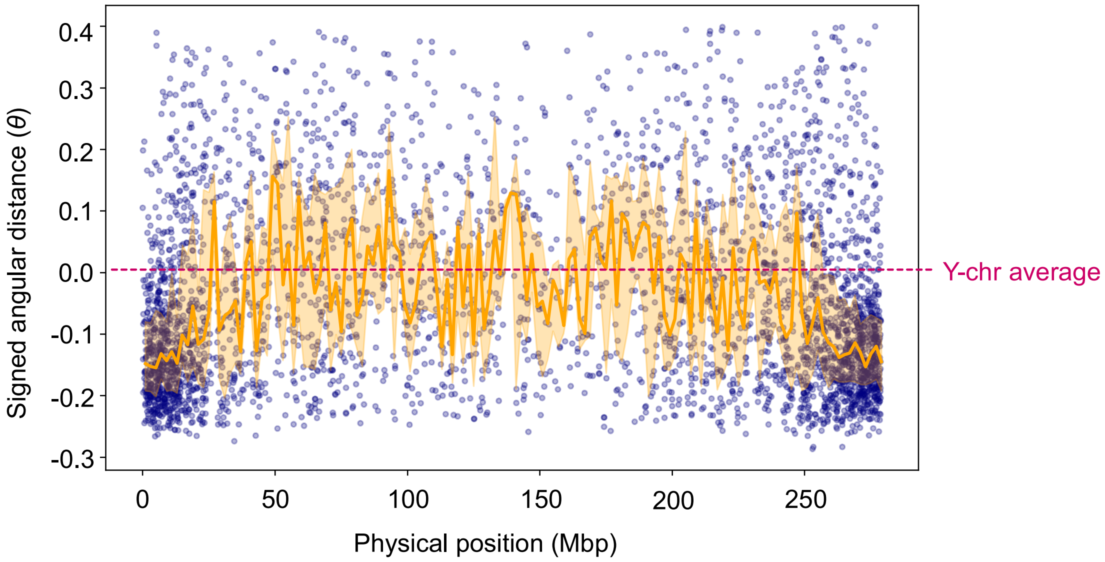


**Supplementary Figure 4 Trajectory of $\theta_\Pi$ along the X chromosome of *Silene latifolia* based on Evo2-derived embeddings**

Consistent with the results obtained using PlantCAD2 (Figure 2c), angular distance analysis based on embeddings generated by Evo2 (7b) showed that $\theta_\Pi$ (angular distance projected onto the X-Y subspace) increases in recombinationally inactive pericentromeric regions, approaching Y chromosome-like properties.

**Supplementary Fig. 5**

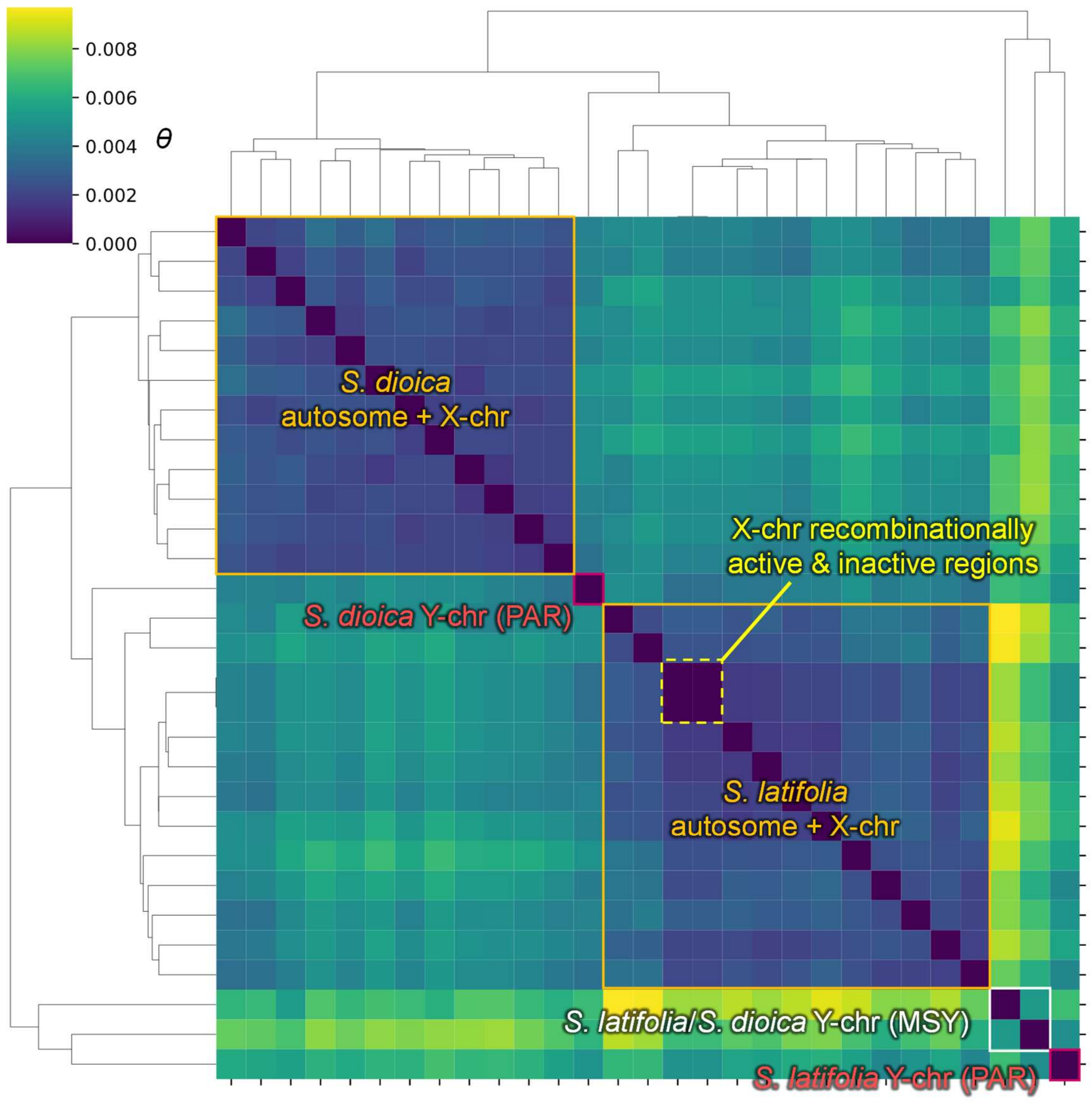


**Supplementary Figure 5 Angular distance analysis with calibration of effects of recombination suppression**

We performed an angular distance analysis by applying an orthogonal projection onto the subspace defined by vectors from recombinationally inactive and recombinationally active regions along the X chromosome of *Silene latifolia*, thereby correcting for (i.e., effectively removing) the influence of recombination suppression. After this correction, the distance between recombinationally active and inactive regions on the X chromosome becomes zero (outlined by dotted yellow lines), and the distance between the Y chromosome and autosomes is also substantially reduced. Nevertheless, the Y chromosome still forms an independent cluster and remains distinguishable from autosomes (outlined by white solid lines). These results suggest that Y chromosome-specific properties cannot be explained solely by recombination suppression. Alternatively, the effects of recombination suppression may be nonlinear, with mutation patterns potentially changing over time depending on the duration of suppression.

**Supplementary Fig. 6**

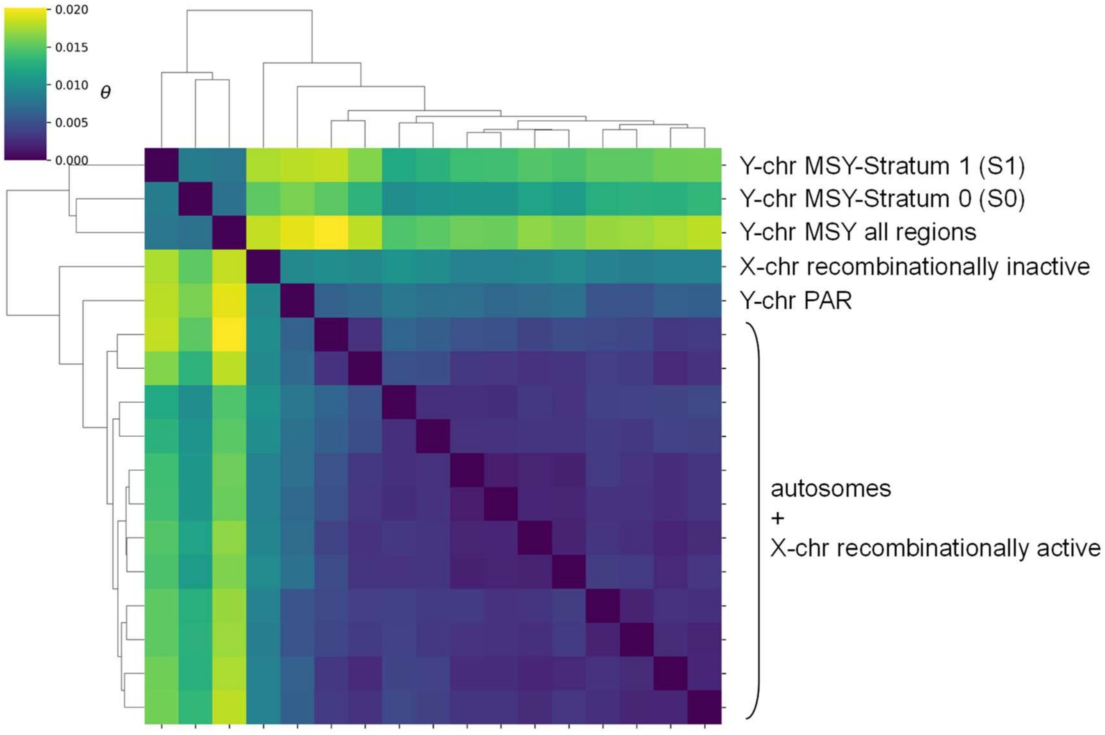


**Supplementary Figure 6 Angular distance analysis of evolutionary strata in the MSY**

In *Silene latifolia*, multiple evolutionary strata reflecting the timing of recombination suppression have been defined. We performed an angular distance analysis in the X-Y subspace for the oldest stratum (S0) and the relatively younger stratum (S1). Both S0 and S1 were included within an independent MSY clade; however, S0 showed a closer relationship to autosomes than S1 (see the heatmap), indicating that the angular distance does not fully reflect the chronological order of recombination suppression.

### Supplementary Fig. 7

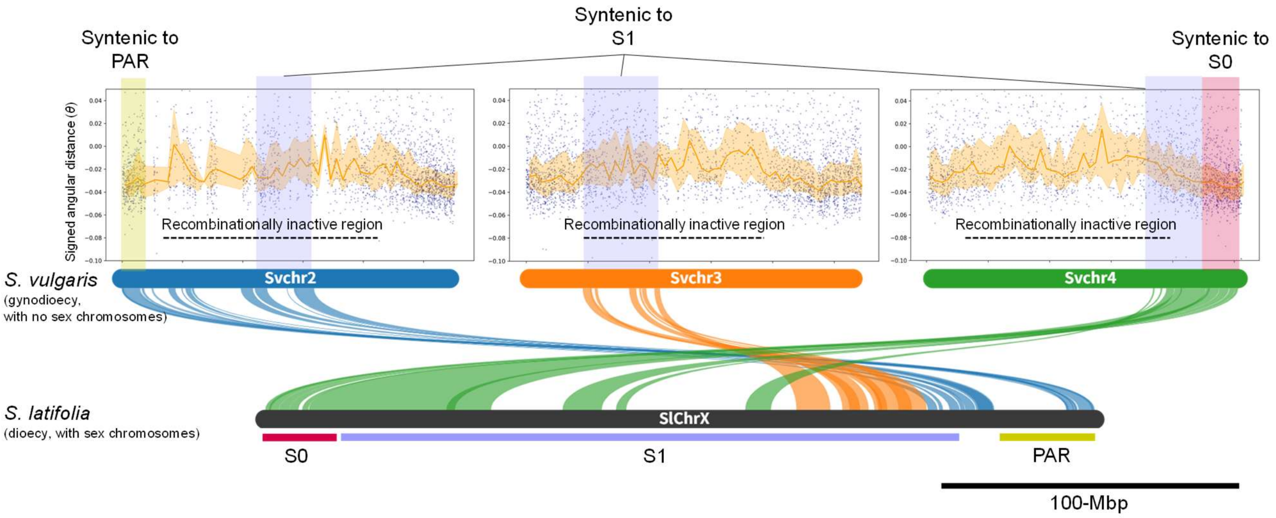


**Supplementary Figure 7 Angular distances in the *Silene* MSY evolutionary strata reflect ancestral characteristics**

We examined syntenic relationships in the closely related gynodioecious species *Silene vulgaris*, which lacks sex chromosomes (Akagi et al. 2025[32]), together with the trajectory of angular distance in the X-Y subspace. The right arm of chromosome 4 in *S. vulgaris*, which is syntenic with the S0 stratum, corresponds to a recombinationally active region and inherently exhibits low $\theta_\Pi$ values. In contrast, most regions syntenic with the S1 stratum (chromosomes 2, 3, and 4 in *S. vulgaris*) correspond to recombinationally inactive regions and are therefore expected to inherently show higher $\theta_\Pi$ values.

**Supplementary Fig. 8**

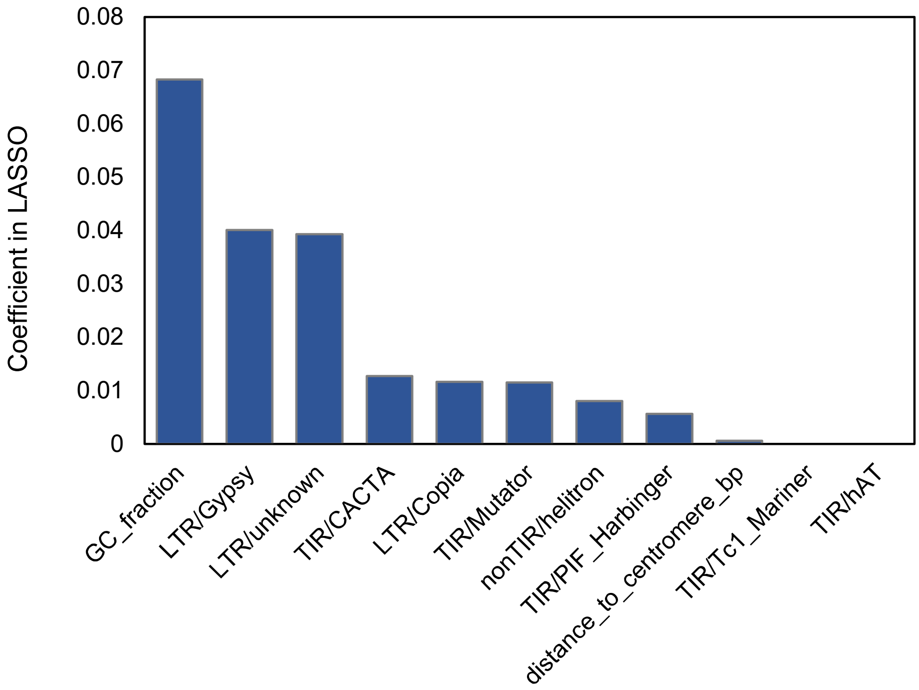


**Supplementary Figure 8 LASSO regression using evo2-derived embeddings**

As in the analysis using PlantCAD2 shown in Figure 2d, we performed LASSO regression on Evo2-derived embeddings, using the angular distance in the X-Y subspace as the response variable. Consistent with the PlantCAD2 results, GC content showed the strongest effect; however, the contributions of transposable elements were generally detected to be higher than with PlantCAD2 embeddings.

**Supplementary Fig. 9**

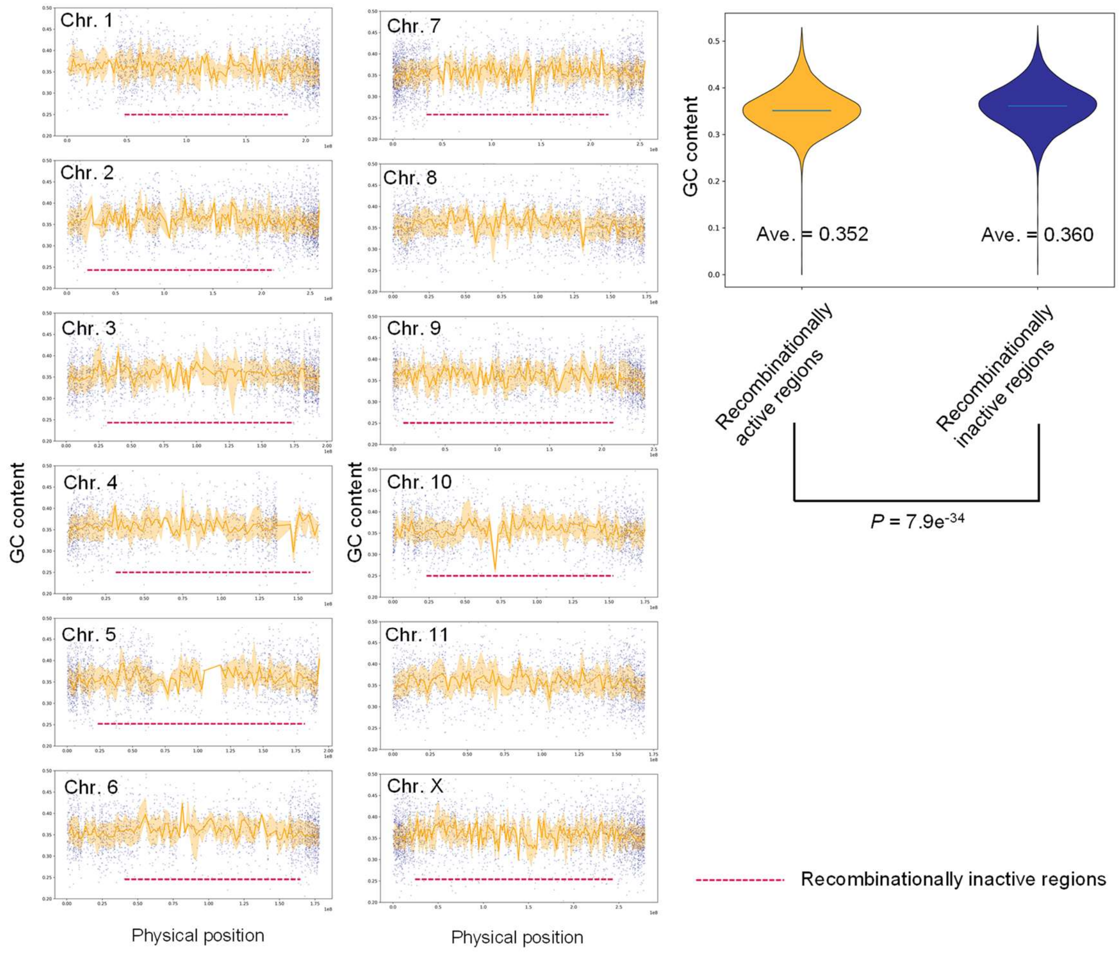


**Supplementary Figure 9 Genome-wide distribution of GC content in *Silene latifolia***
We show the genome-wide variation in GC content across the same set of regions used for embedding generation. GC content tends to increase in recombinationally inactive regions (indicated by magenta dashed lines), particularly in pericentromeric regions, albeit not prominent. This trend is also reflected in the violin plots (right panel), which include medians and show a significant difference between recombinationally inactive and active regions ($p$ = 7.9 x $10^{-34}$, two-sided Student's *t*-test). Chromosomes 8 and 11 were excluded from the analysis because detailed genetic maps are not available[32].

**Supplementary Fig. 10**

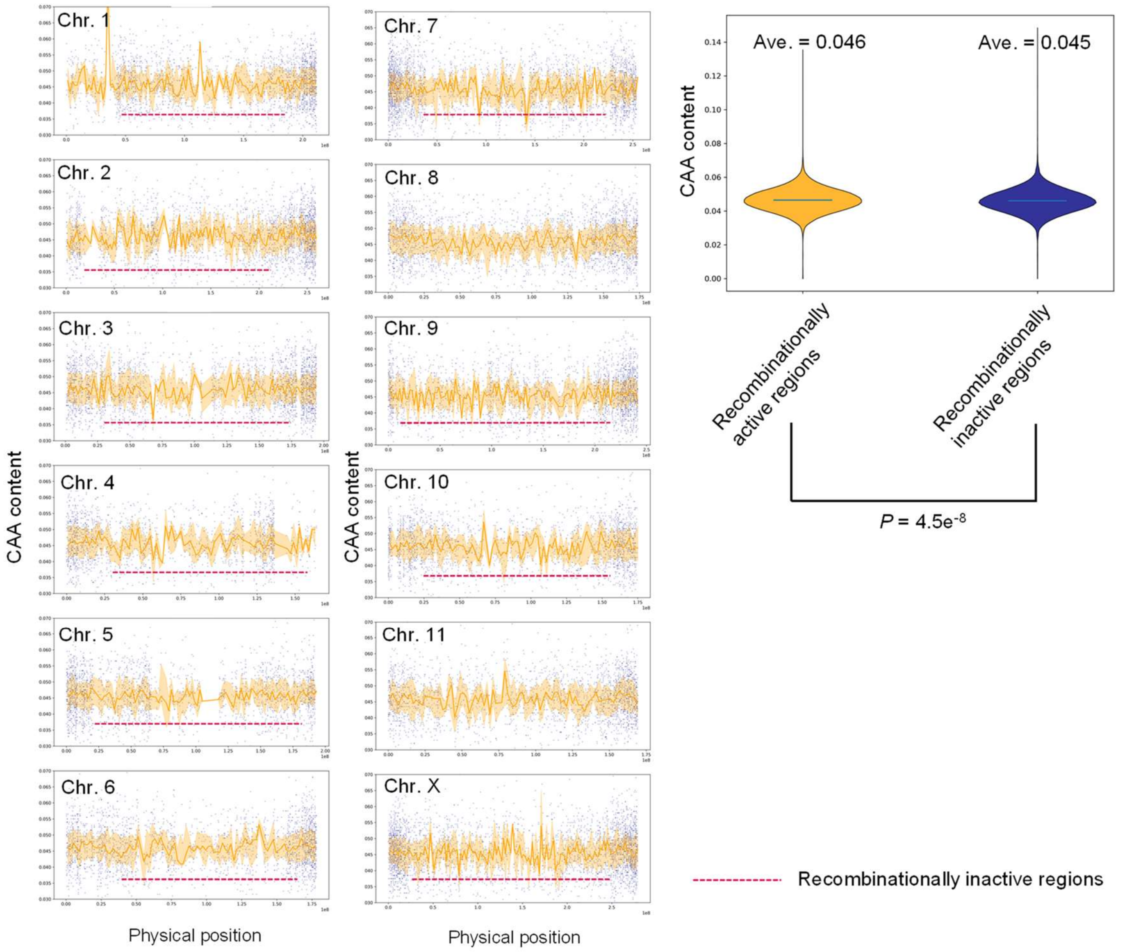


**Supplementary Figure 10 Genome-wide distribution of CAA content in *Silene latifolia***

We show the genome-wide variation in CAA content across the same set of regions used for embedding generation. In inverse proportion to GC content, CAA content tends to decrease in recombinationally inactive regions (indicated by magenta dashed lines), although this trend is further inconspicuous. The significant difference could be observed between recombinationally inactive and active regions ($p$ = 7.9 x $10^{-34}$, two-sided Student's *t*-test), as given in the violin plots (right panel) which include medians. Chromosomes 8 and 11 were excluded from the analysis because detailed genetic maps are not available[32].

**Supplementary Fig. 11**

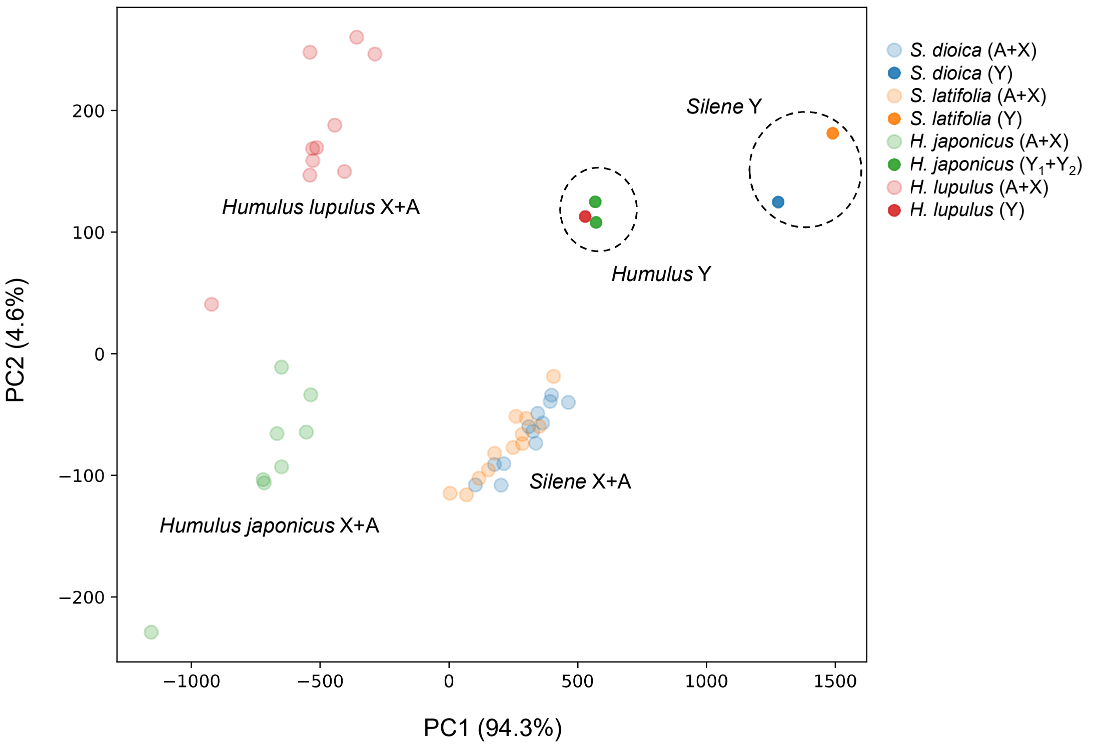


**Supplementary Figure 11 Separation of Y chromosome-specific components by PCA using evo2-derived embeddings**

We show the distribution along principal component (PC) axes that exhibit statistically significant differences between autosomes/X chromosomes and Y chromosomes. PC1 primarily contributes to the separation between the genera *Silene* and *Humulus*, while PC2 contributes to species-level separation within *Humulus*. For Y chromosome separation, Y-specific clusters were detected as in the PlantCAD2 analysis; however, the direction and norm (or distance) of the Y chromosome group relative to autosomes were not completely consistent between *Silene* and *Humulus*.

**Supplementary Fig. 12**

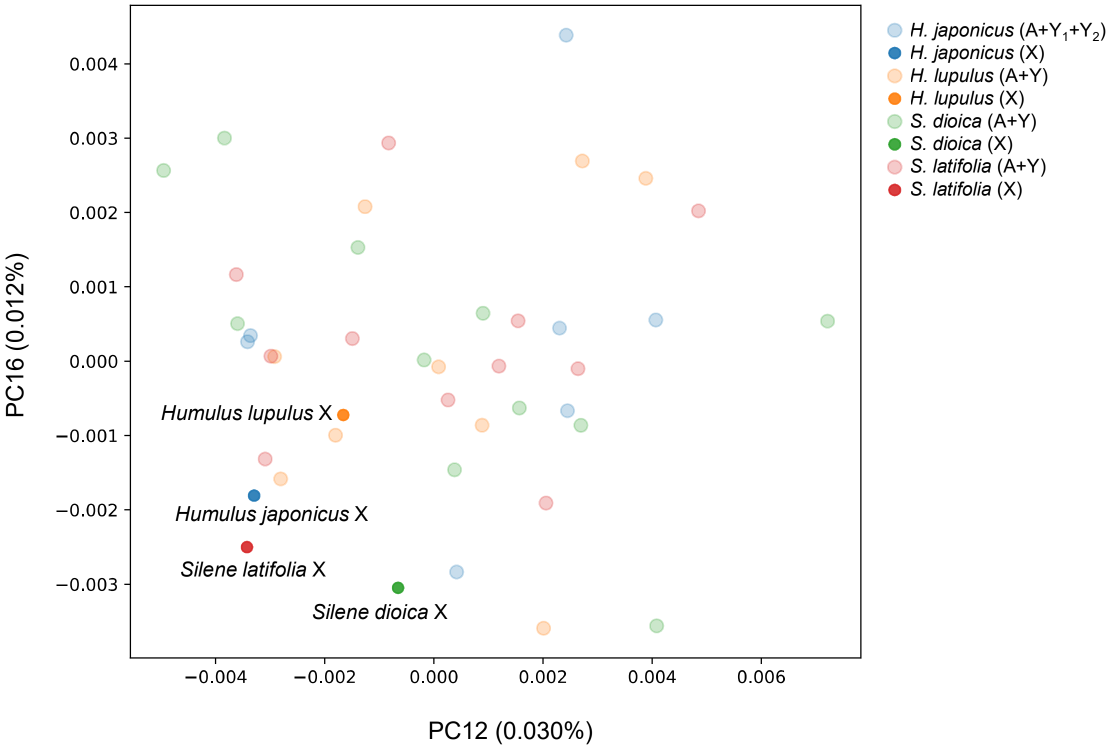


**Supplementary Figure 12 Exploration of X chromosome-specific principal components from PlantCAD2-derived embeddings**

We performed PCA on PlantCAD2-derived embeddings and extracted two principal components (PC12 and PC16) that showed significant differences ($p < 0.05$) between the X chromosome group and the combined autosome and Y chromosome group. The distributions of each chromosome were then plotted. Although the X chromosome group showed a tendency to separate from the others, the boundary was ambiguous and did not form a completely distinct cluster. Note that we also tested embeddings with motif-level signal enhancement using GeM pooling, but no clear X-specific separation was observed.

**Supplementary Fig. 13**

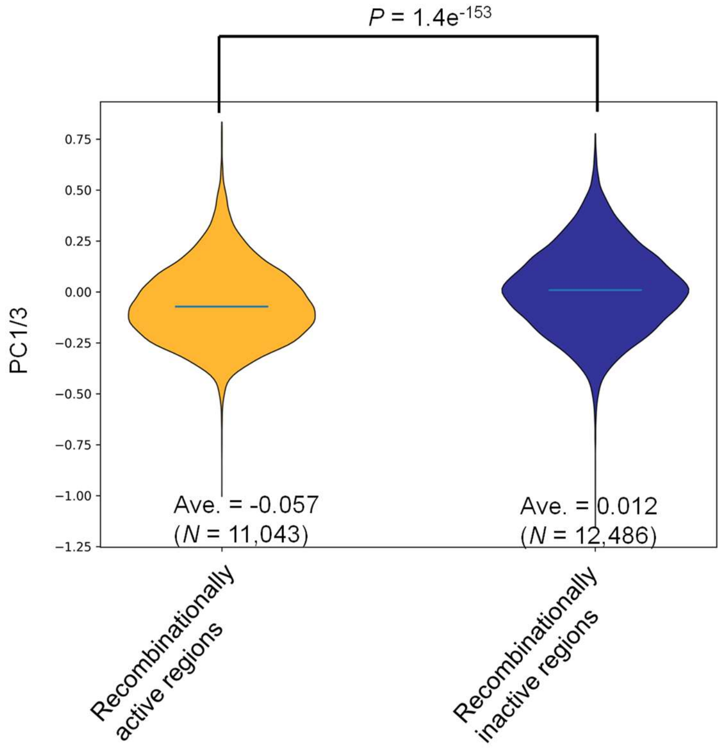


**Supplementary Figure 13 Distribution of PC1/PC3 values in genome-wide recombinationally inactive and active regions in *Silene latifolia***

We computed PlantCAD2-derived PC1/3 values for all gene promoter regions in *Silene latifolia* and compared their distributions between recombinationally inactive and active regions using violin plots (including medians). PC1/3 values were significantly elevated in recombinationally inactive regions ($p$ = 1.5 x $10^{-143}$ in two-sided Student's *t*-test), suggesting a shift toward Y chromosome-like properties. However, the median PC1/3 value in recombinationally inactive regions remained significantly lower than that of the Y chromosome ($p$ = 2.3 x $10^{-54}$ in two-sided Student's *t*-test).

**Supplementary Fig. 14**

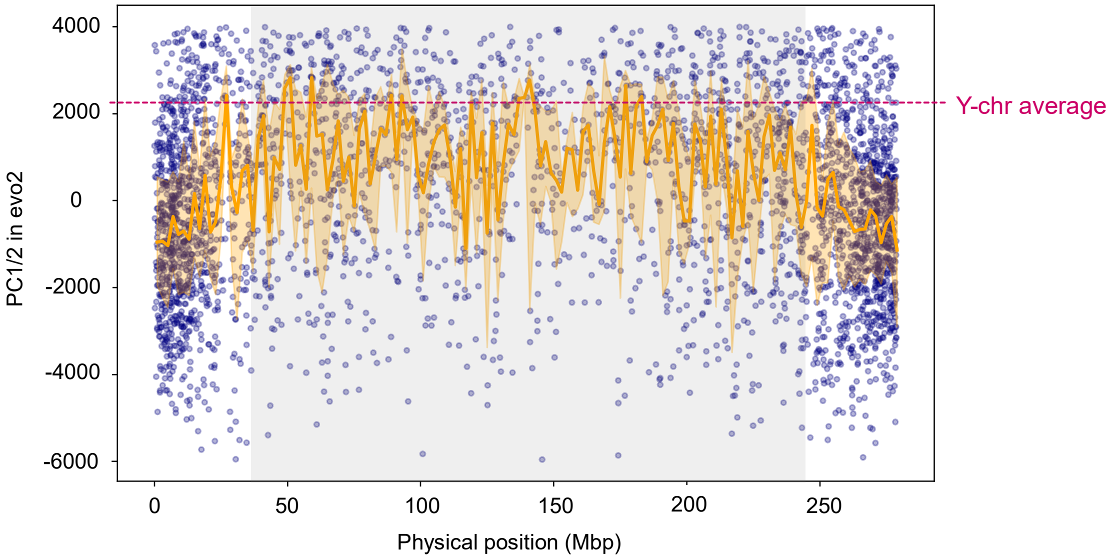


**Supplementary Figure 14 Trajectory of PC1/2 values along the *Silene latifolia* X chromosome derived from evo2 embeddings**

Consistent with the results based on PlantCAD2-derived data (Figure 3b), PC1/2 values tend to increase in recombinationally inactive regions (highlighted in gray) along the X chromosome, suggesting a shift toward Y chromosome-like properties. Values in pericentromeric regions are closer to the mean values of the Y chromosome, than with the PlantCad2-derived data (Figure 3b). This suggests that these combined principal components in Evo2 may capture sequence contexts immediately arising from recombination suppression, regardless of its duration.

**Supplementary Fig. 15**

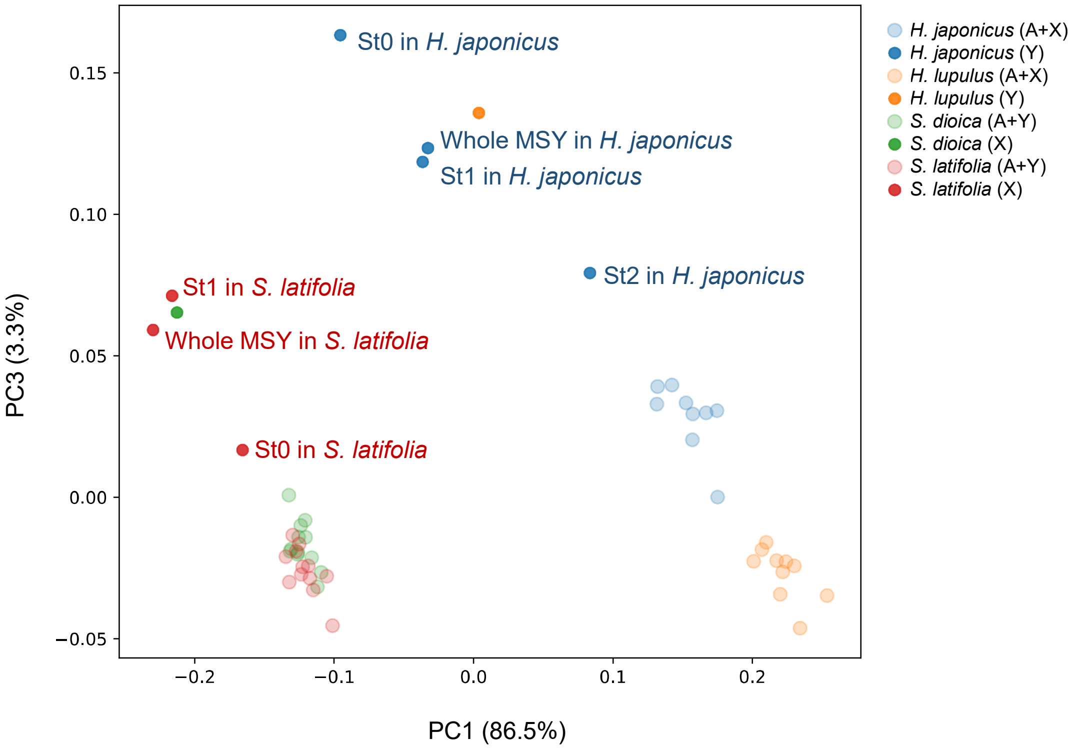


**Supplementary Figure 15 Distribution of PlantCAD2-derived PC1/3 values across evolutionary strata within the MSY**

In the PC1/PC3 space shown in Figure 3a, we additionally plotted the distributions of S0 and S1 in *Silene latifolia*[32], as well as S0, S1, and S2 in *Humulus japonicus*[34]. In *H. japonicus*, the divergence of each stratum from autosomes was consistent with the relative ages of stratum formation: the oldest stratum (S0) was the most distant from autosomes, whereas the relatively younger stratum (S2) showed a distribution closer to autosomes. In contrast, in *S. latifolia*, the distance from autosomes did not correspond to the relative ages of the strata; the relatively younger S1 was located farther from the autosome cluster than S0. As discussed in Supplementary Fig. 7, this pattern likely reflects the ancestral properties of the chromosomal regions from which S0 and S1 originated.
